# Unique and assay specific features of NOMe-, ATAC- and DNase I-seq data

**DOI:** 10.1101/547596

**Authors:** Karl JV Nordström, Florian Schmidt, Nina Gasparoni, Abdulrahman Salhab, Gilles Gasparoni, Kathrin Kattler, Fabian Müller, Peter Ebert, Ivan G. Costa, DEEP consortium, Nico Pfeifer, Thomas Lengauer, Marcel H Schulz, Jörn Walter

**Author notes:** These authors contributed equally to this work (Florian Schmidt, Nina Gasparoni, Abdulrahman Salhab, Jörn Walter). Department of Genetics, Stanford University School of Medicine, Stanford, CA 94305, USA. Methods in Medical Informatics, Department of Computer Science, University of Tübingen, Tübingen, Germany (Nico Pfeifer). Institute for Cardiovascular Regeneration, Goethe University Frankfurt and German Center for Cardiovascular Research, Partner site Rhein-Main, Frankfurt am Main 60590, Germany (Marcel H Schulz). http://www.deutsches-epigenom-programm.de/ (DEEP consortium).

## Abstract

Chromatin accessibility maps are important for the functional interpretation of the genome. Here, we systematically analysed assay specific differences between DNase I-Seq, ATAC-Seq and NOMe-Seq in a side by side experimental and bioinformatic setup. We observe that most prominent nucleosome depleted regions (NDRs, e.g. in promoters) are roboustly called by all three or at least two assays. However we also find a high proportion of assay specific NDRs that are often “called” by only one of the assays. We show evidence that these assay specific NDRs are indeed genuine open chromatin sites and contribute important information for accurate gene expression prediction. While technically ATAC-Seq and DNAse I-Seq provide a high NDR calling rate for relatively low sequencing costs in comparison to NOMe-Seq, NOMe-Seq singles out as it provides a multitude of information: it allows to not only detect NDRs but also endogenous DNA methylation, genome wide segmentation into heterochromatic A/B domains and local phasing of nucleosomes outside of NDRs. In summary our comparison strongly suggest to consider assay specific differences for the experimental desgin and for generalized and comparative functional interpretations.

## INTRODUCTION

The eukaryotic genome is largely organized in nucleosomes - the basic unit of chromatin. The precise mapping of nucleosome occupancy and the accessible DNA provides a widely used molecular approach to monitor chromatin organization in cells and the specific and local impact on gene expression. Three main experimental approaches, DNase I-Seq, ATAC-Seq and NOMe-Seq, are mainly used to comprehensively analyze chromatin accessibility on a genome wide level. All these approaches have in the meantime been scaled down to single cell analyses making them the prime assays for functional studies (1–5). In some cases these assays have even be applied to simultaneously measure gene expression (RNA-Seq) and chromatin accessibility (NOMe-Seq, ATAC-seq) from the same cell (5–7).

Several independent studies demonstrated that each of these methods come with advantages and disadvantages but so far very few systematic comparative analyses have been performed using standardized procedures. Our study fills this gap aiming to evaluate the comparability and complementarity of the three methods and at the same time showing their individual advantages or limitations to identify and interpret epigenetic and chromatin data.

The oldest and still widely used method to analyze nucleosome occupancy and positioning is DNase I-Seq (8) (see Figure 1). DNase I cuts at freely accessible DNA to release small DNA fragments, which when coupled to next-generation sequencing (NGS) can be traced back to the genome to identify open chromatin regions. Extensive DNase I studies by ENCODE show that this approach boosts the identification of regulatory elements in different cells and showed that numerous genetic variants identified in genome-wide association studies (GWAS) locate to such DNase I hypersensitive regions (9). Still, the general use of DNase I-Seq for open chromatin mapping can be quite difficult. First of all, DNase I experiments usually require a fair amount of pre-testing to identify the right incubation conditions, because, among other factors, under-or over-digestion of the chromatin will greatly influence the detection rate of open chromatin sites (10). For this the abundance of primary cell material can often be a problem. Usually several experiments with different amounts of DNase I have to be performed to determine the optimum conditions. Once conditions are established the method can be scaled down to even single cells as showed by (2). Using paired end sequencing the prediction of nucleosome depleted regions (NDRs) from aligned DNase I-seq reads can be performed with conventional peak callers developed for ChIP-Seq such as MACS2 (11).

**Figure 1.**
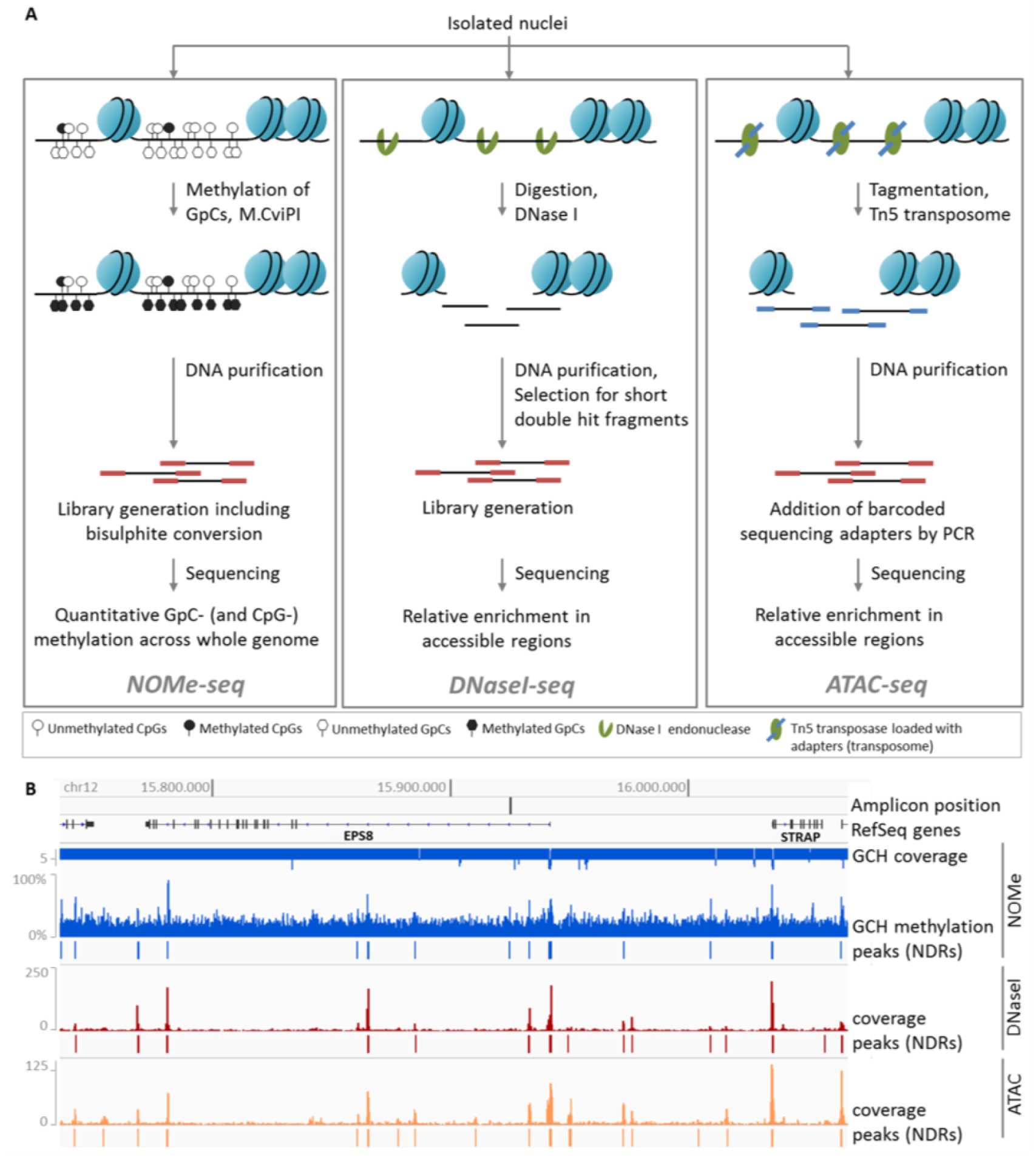
Chromatin accessibility assays. A. NOMe-seq (left): Isolated nuclei are treated with M.CviPI, which methylates cytosines in GpC-dinucleotides. The DNA is purified, subjected to library preparation including bisulfite-conversion and sequenced. The resulting data give quantitative measures of GpC-and endogenous CpG-methylation at base-pair resolution across the whole genome. DNase I-seq (middle): After isolation of nuclei, chromatin is digested with DNase I. The DNA is purified and short double hit fragments are selected, which are used for library preparation and sequencing. The obtained data represent a relative enrichment of double-hit fragments in accessible regions. ATAC-seq (right): Isolated nuclei are treated with Tn5 transposase, which is loaded with sequencing adapters. After the tagmentation of chromatin (fragmentation and tagging by the transposome), the DNA is purified, barcoded by PCR and sequenced. The data give a relative read-out of tagmented fragments across accessible regions. B. Snapshot of chromatin accessibility data relative to genomic coordinates for section of chr12. NOMe-seq (blue): Quantitative measure of GCH-methylation and NDRs called with gNOMeHMM are shown. The GCH coverage track indicates if number of reads is 5 (downward bars) or higher (upward bars). DNase I-(red) and ATAC-seq (orange): Read coverage and NDRs called with MACS2 are shown.

A second method with growing popularity is ATAC-Seq (12) (see Figure 1), which stands for assay for transposase-accessible chromatin. ATAC-Seq uses hyperactive Tn5 transposase-mediated cutting of genomic DNA combined with ligation-mediated insertion of DNA oligonucleotides which are pre-loaded (in vitro) to the enzyme. Following DNA isolation and PCR-amplification, libraries can directly be used for NGS. Because of its relative technical simplicity and high sensitivity this method is increasingly used (13–17). ATAC-Seq requires very low amount (down to single cells) of (even frozen) input material to generate comprehensive maps (1,000 50,000 cells). Since the ATAC reaction is an “end-point” reaction it reduces the risk of chromatin over-digestion. Additionally, due to the relative simplicity of the protocol, ATAC-seq allows for high-throughput application to hundreds of bulk samples at relatively low cost making it applicable to larger clinical cohorts (18). A drawback of ATAC-seq is a frequently observed enrichment of mitochondrial (MT) sequence reads due to the unprotected nature of MT DNA which appears to be a preferred target for the Tn5 transposase in the cell. This can be avoided by depleting unwanted reads with a CRISPR/Cas9 based strategy (19, 20). This addition however makes the method more laborious. ATAC-Seq data can be processed using standard peak peak callers developed for ChIP-Seq such as MACS2 (11).

The third method with growing popularity is a nuclease free method called NOMe-Seq which stands for “nucleosome occupancy and methylation”. NOMe-Seq utilizes the enzyme M.CviPI that specifically methylates cytosine dyads in a GpC sequence context originally used to identify local open chromatin regions (21). Following the incubation of permeabilized cells or nuclei with M.CviPI, the extracted DNA is subjected to conventional bisulfite conversion followed by either regional (targeted) or genome-wide sequencing (WGBS). With this approach the DNA-methylation levels at GpC sites (NOMe reaction) and at endogenous CpG sites are determined simultaneously. The GpC methylation can be used to map DNA accessibility and determine nucleosome free regions. NOMe also has some interesting (partially unique) technical features: i) NOMe-Seq only requires low amounts of input material (down to single cells) and is applicable for many cell types. ii) The in vitro methylation step is an endpoint reaction reducing the risk of over-exposure. iii) NOMe-Seq data provide a direct and single molecule quantitative readout, measuring the accessibility for each GpC in a single chromosome. iv) NOMe can be seamlessly adopted for region-specific or whole genome-wide analysis of open chromatin. v) NOMe-seq delivers the endogenous CpG methylation as a second readout enabling the simultaneous analysis of chromatin accessibility and DNA methylation on the same molecule in one experiment. The NOMe-Seq “readout” of open chromatin sites is limited to GpC containing sequences and requires a high sequencing depth.

Recent NOMe-based assays, scNMT-seq allows the combined sequencing of nucleosome occupancy together with transcriptome of the same cell (5). The current methods for NOMe-seq data interpretation leave room for improvement. (22) recently developed the findNDR tool, which computes enrichment scores of methylated reads for fixed size windows against a genome-wide average methylation rate. Here, we present a new algorithm for detecting NDRs in NOMe-seq data that allows for variable size length, performs a proper FDR correction, corrects for local, regional methylation backgrounds and corrects for abundance artifacts due to amplifications as commonly found in cancer genomes.

To perform a direct comparison of DNase I-, NOMe-and ATAC-Seq data we decided to produce all data from a standard model cell line HepG2. We performed all experiments in our lab using the same stock of cells and the same cultivation conditions to minimize technical confounding variables. We deliberately used bulk cells to obtain deep and comprehensive genome wide overviews for each method. We observe that all three methods have a considerable overlap in detecting (strong) major open chromatin regions but also deviate from each to a large extend in other regions. We find that none of the methods calls the entire spectrum of open chromatin sites. This becomes most apparent when using these data for functional prediction of gene expression. We believe that our detailed comparison will contribute to better understand the technical and experimental limits of each of the three methods and at the same time allow to rationalize which approach should be used to address particular questions.

## METHODS

### Sequencing, pre-processing

Fastq files were trimmed for adapter sequence and low quality tails (Q*<* 20) with trim galore (23), after which they were mapped to the human (hs37,1000G) genome (24). For WGBS and NOMe data, this was done with *GSNAP* (25) and for DNase I and ATAC data, *GEM* (26) was utilized (See Table S1).

### WGBS and NOME

Unmapped reads were removed with *samtools* (27), before further processing with the Bis-SNP pipeline (28). Initially, reads were remapped in regions close to known insertions or deletions as supplied by Database of Single Nucleotide Polymorphisms (dbSNP). Bethesda (MD): National Center for Biotechnology Information, National Library of Medicine. (dbSNP Build ID: 138). Available from: http://www.ncbi.nlm.nih.gov/SNP/. Duplicated reads were marked with Picard tools (http://broadinstitute.github.io/picard) and potentially overlapping sections between two paired reads were clipped with bamUtils (29). The quality was recalibrated in the context of dbSNP. Finally, methylation levels were called for all cytosines, and extracted with a modified version of the Bis-SNP *vcf2bed.pl* helper-script. For the NOMe samples, bed files were generated for cytosines in GCH and HCG context(, where H is the IUPAC code for A, C or T). This corresponds to artificial and natural methylation, respectively. For WGBS samples, files for CG context were generated.

### DNase I and ATAC

Duplicated reads were annotated with Picard tools. MACS2 (30) was used to call NDRs. In comparison to ChIP-seq, the cutting with DNase I and inclusion of adapters with the transposase in ATAC puts the focus on the start and end of a fragment, therefore MACS was executed with the following parameters: `--shift -100`, `--extsize 200`, `--nomodel` and `--keep-dup all`. All duplications were kept as MACS only takes one end of the fragment into account when designating duplicated reads compared to Picard tools that makes use of both.

### External data

External data sources are listed in data file

### Finding open chromatin regions with NOMe data

Given methylation values *M*_*i*_,*i* = 1,*…,m*, for *m* cytosines in GCH context in the human genome, we use a Hidden Markov Model (HMM) to segment all cytosines in GCH context into two states (i) NDR and (ii) occupied region. As the methylation values are in range [0,1] we use a binomial distribution in each HMM state. We use the Baum-Welch algorithm to fit the 2-state HMM using the *HiddenMarkov* R package (31). We stop the parameter optimization after either 1000 iterations or if the likelihood between two consecutive rounds drops below 10*-*3. We fit an HMM for each chromosome and use parallelization with the SNOW package for fast computation. Each GC nucleotide is predicted to be open or closed based on posterior decoding using the fitted binomial HMM. Stretches of predicted NDRs, so called peaks, are further ranked by significance.

To assign significance to the potential open chromatin regions found by the HMM, we computed empirical false discovery rates (FDRs) as well as the corresponding q-values. (32) The false discovery rate at significance threshold *t*(*FDR*(*t*)) is the expected value of the proportion of false discoveries significant at threshold *t*(*f* (*t*)) divided by the total number of discoveries that are significant at threshold *t*(*s*(*t*)):

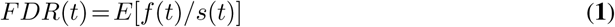

This can be approximated by *E*[*f* (*t*)]*/E*[*s*(*t*)]. Using the HMM on our data set, we could directly get an estimate for *E*[*f* (*t*)] by counting regions with a p-value smaller than or equal to *t*. To estimate *E*[*s*(*t*)], we generated null data from the real data by shuffling methylation levels of the input data, leaving all other parts of the data intact. We then computed the segmentation for this data. This was performed to get the p-value distribution of regions falsely labeled as open.

P-values were calculated with a one-sided Fisher’s exact test, contrasting the number of observations of methylated and unmethylated cytosines in GCH context within the region to a that of a background region. The background was selected as the closest 4kb of closed chromatin up-and down-stream of the tested region. Due to the coverage dependence of the chosen test, we implemented an automatic stratification step based on non-parametric mixture model clustering with the *Mclust* R-package (33). Assuming that regions with deviating copy numbers is the exception, we let the median represent the common coverage, and optimize the shrinkage parameter to minimize the number of clusters with mean coverage below the median. After each loci are assigned to a cluster, FDR values are estimated within each cluster. The gNOMeHMM package can be retrieved from https://github.com/karl616/gNOMePeaks.

### Processing of RNA-seq data

*TopHat 2.0.11* (34) and *Bowtie 2.2.1* (35) were used to generate BAM files of RNA-seq reads, for NCBI build 37.1 in `--library-type fr-firststrand` and `--b2-very-sensitive` setting. Gene expression was quantified using *Cufflinks 2.0.2* (36), the hg19 reference genome using the options `frag-bias-correct`, `multi-read-correct`, and `compatible-hits-norm`.

### Determine characteristics of DNase I, ATAC, and NOMe NDRs

To better understand the characteristics of the DNase I endonuclease, the Tn5 transposase, and the GpC methyltransferase M.CviPI, we generated sequence motifs, DNA shape predictions, as well as investigated DNA methylation at DNase I-seq and ATAC-seq cut-sites and at GpC sites for NOMe, respectively. We obtained the considered sequences using the *bedtools getfasta* (37) command on the 5’-cut-sites/GpC sites retrieved from the aligned reads. Note that all ATAC-seq sequences are shifted by 4bp upstream, to consider the center of the Tn5 transposase (12). Per GpC sites, we sampled reads according to the methylation state of the respective GpC.

#### Generation of sequence motifs

Using 59,850,858 DNase I-sequences, 30,108,148 ATAC-seq sequences, and 126,202,679 NOMe-sequences, we generated sequence logos using the *ggseqlogo* R-package (38). The sequence motifs reported in literature are 6bp for DNase I (39) and 20bp for ATAC (12). To our knowledge, no bias motif has been reported in literature for NOMe-seq. To ensure these are captured and to harmonize with other figures, we used 31bp centered on the enzyme activity sites of each assay.

#### Shape prediction

We use the *DNAshapeR* R-package (40) predictions for the minor groove width (MGW), Roll, propeller twist (ProT), and helix twist (HelT). Due to memory limitations, we randomly selected 2 million sequences per assay, constructed as described above, to be used for the shape computations. All spatial features are computed for each assay in a 31bp window centered at the enzyme activity site.

#### DNA methylation

For each assay and sample, CpG methylation was extracted in 40bp windows centered on an enzyme event (bedtools (37)). Average methylation, weighted to coverage, was calculated for each relative position.

### Logistic regression classifier for assay-specific NDRs

To identify characteristics of NDRs identified with only one distinct assay, we learn a multi-class logistic regression classifier using a variety of sequence based features, explained in the next section.

#### Feature definition

Within each assay specific NDR, we computed

- A, T, C, and G content,
- CG content,
- CpG and GpC count,
- average CpG methylation,
- NOMe-seq coverage,
- and counts for all 5-mers.

We excluded GpC methylation as it would be an obviously strong feature for NOMe-seq NDRs, potentially overshadowing the remaining features. Additionally, we excluded NDR length as a feature, as this would mainly resemble peak caller specificities, as shown before for different peak callers on DNase I-seq data (11). The aforementioned features are computed for 12,415 DNase I NDRs, 19,323 ATAC-seq NDRs, and 19,453 NOMe NDRs (excluding the Y-chromosome).

#### Logistic regression

We learn a multiclass logistic regression classifier using elastic net regularization, as implemented in the *glmnet* R-package (41). Elastic net regularization is producing sparse models and at the same times distributes the regression coefficients among correlated, yet predictive features, a property known as the grouping effect. This is achieved by combining two penalty terms, the lasso(L1) and the ridge(L2) penalty:

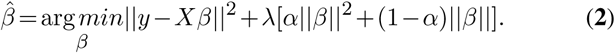

Here, 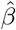 denotes the estimated model coefficients, *β* are the model coefficients, *X* is the feature matrix, which is scaled and centered, *y* is the response vector containing class assignments and *α* is a parameter regulating the trade-off between *L1* and *L2* penalty. We optimize the *α* parameter in a grid search from 0.0 to 1.0 with a step-size of 0.01. This is performed in scope of a 10-fold Monte-Carlo cross-validation procedure, in which the data is randomly split into 80% training and 20% test data, assuring that both sets are balanced (see Figure S1). Within each fold, we perform a nested-six fold inner cross validation to fit regression coefficients and determine the *λ* parameter controlling the total amount of regularization. We choose *λ* over the inner folds according to the minimum miss-classification error (*λ*_*min*_). Final model coefficients are determined using *λ*_*min*_ on the entire training data set. Model performance is computed in terms of accuracy (ACC) on balanced hold-out test data using a 3*x*3 confusion matrix *C*:

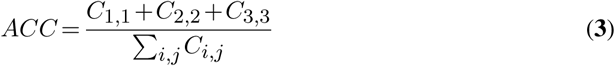

Note that, since we perform a 3-class classification, randomness is reflected by *ACC ≤* 0.33.

### Computation of nucleosome distances

We predicted nucleosome positions based on ChIP-seq data by running *Nuchunter* (42) with default parameters. The distance between neighboring predicted nucleosomes was calculated, the 220,315 gaps between 0 and 500bp were selected and stratified into bins of 50bp. Overlapping these gaps with NDRs and transcription factors allowed us to calculate enrichments in different size regimes. The enrichment was calculated as the difference between the observed and estimated number of overlapping features. For each feature and size bin, the estimated number of overlaps was calculated by distributing the observed overlaps over the bins in proportion to the number of expected overlaps. For plotting, the absolute values were log transformed, while keeping the sign of the original value

### NDR clustering

For each set of NDRs, we calculated the average NOMe signal intensity in tiled 10bp bins spanning 1kb up-and down-stream relative to the NDR summit. Subsequently, we identified all combinations of overlapping regions between the three NDR sets and the signal from these were combined. In the absence of a NDR the loci were defined by another assay, prioritized as NOMe, DNase I or ATAC. That is, a lacking NOMe NDR was approximated by DNase I and only if both NOMe and DNase I lacked NDR, the ATAC NDR was used as substitute. The resulting matrix consisted of rows with 200 bins from each assay. It was clustered into fifteen clusters with the k-means algorithm.

The resulting clusters were characterized with respect to the fraction of NDRs overlapping a ChromHMM segmentation of the DEEP HepG2 histone data and transcription factors with LOLA and an extended version of its core data base (43). The ChromHMM states are labeled in agreement to the 15-state core segmentation provided by Roadmap (44).

### Linear regression predicting gene expression

To learn about the relationship between chromatin accessibility and gene expression, we fitted linear regression models predicting gene expression from predicted TFBSs.

#### Feature definition

We compute TFBS features using *TEPIC* (45, 46) in all ATAC 𝒜, DNase I *𝒟*, and NOMe *𝒩* NDR sets. Additionally, we consider the intersection *ℐ* of the three sets, as well as their union *𝒰*:

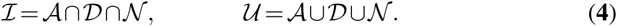

Furthermore, we consider three NDR sets extending 𝒜, 𝒟, and 𝒩 to match | *𝒰*|by randomly sampled regions that do not overlap with any of the already included open chromatin NDRs. We refer to these NDR sets with 𝒜_*R*_, 𝒟_*R*_, and 𝒩_*R*_ respectively. Thus, we consider in total eight different NDR sets 𝒫_*j*_ for *j* = 1*…*8.

For each NDR *p ∈𝒫*_*j*_, we compute TF affinities *a*_*p,t*_ for TF *t* using *TRAP* (47), normalized according to the length of the respective NDR |*p*|, for a set of 726 TFs. Position weight matrices are taken from the *TEPIC 2.0* repository (46).

TF affinities *a*_*p,t*_ are combined to TF-gene scores for gene *g* as suggested in (45),

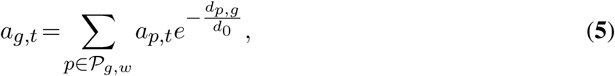

using a windows size *w*=50kb. In addition to TFBS features, we consider NDR length *l*_*g*_ and NDR count *c*_*g*_ as additional features per gene (as introduced in (48)):

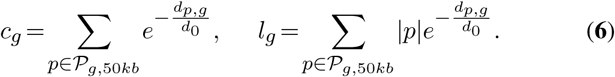

Values for *a*_*g,t*_, *c*_*g*_, and *l*_*g*_, are computed for all protein-coding genes that are associated to at least one DHS site. Where *𝒫*_*g*,50*kb*_ denotes all peaks in a 50kb window around gene *g*.

#### Linear regression

As for the logistic regression, we use elastic net regularization to learn a linear model of gene expression. Here, a row of the feature matrix *X* is composed of the TF gene scores *a*_*g,t*_ as well as the values of *c*_*g*_ and *l*_*g*_ for a distinct gene *g*. The actual gene expression is denoted by the response vector *y*.

The learning strategy is identical to the one explained above, with the difference that the cross-validation error is measured as mean-squared error instead of miss-classification error. Final model performance is assessed using Spearman correlation computed between the actual gene expression *y*, and the predicted gene expression *y* ŷ. Models are learned independently for each NDR set.

### Hi-C data

Hi-C data of HepG2 from (49) was used (GSE113405). The interaction matrix and TADs were generated as described before (49). A/B compartment predictions were generated using HOMER tools (50) with the *runHiCpca.pl* command with the following parameters: `-res 25000 -window 50000`. Positive values of pca1 correspond to A compartments while the negative values correspond to the B compartments. The compaction score Distal-to-Local [log2] Ratio (DLR) was calculated using *analyzeHiC* command with the following parameters: `-res 5000 -window 15000 -compactionStats auto`. Figure 2A was generated using pyGenomicTracks (51).

**Figure 2.**
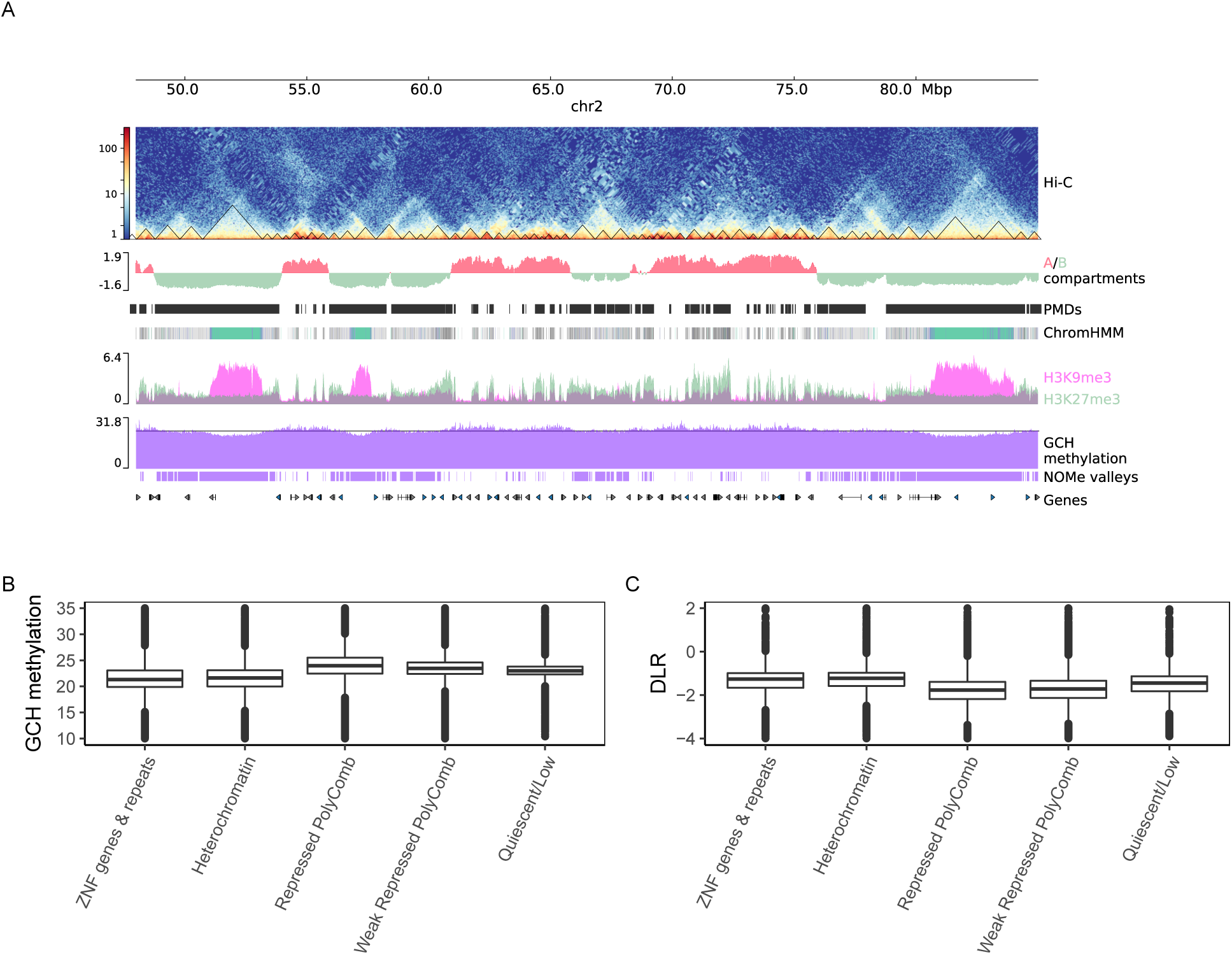
Genome wide chromatin landscape is reflected by NOMe signal. A) zoomed out view (chr2:48,000,000-85,000,000) of open chromatin assay (NOMe-Seq) with different epigenomic data tracks: from top to bottom; Hi-C contact matrix of HepG2 with called TADs (triangles), A/B compartments, ChromHMM segmentation using six histone marks (see Methods), two overlapping heterochromatic marks H3K9me3 and H3K27me3, GCH methylation signal filtered for NDRs, large-scale segmentation of NOMe signal (the horizontal line represents the value chosen by the classification model to define valleys) and UCSC genes. Valleys are identified from NOMe (see Methods) coincide with B-compartments, PMDs from (49), repressed polycomb (gray) and heterochromatic domains (Turquoise) and mostly pronounced at ZNF/repeats regions (Medium Aquamarine). B) GCH methylation levels are low at heterochromatic and ZNF/Repeats regions, while they are high in the repressive polycomb regions. C) Compactness score derived from Hi-C data (DLR, see Methods) is higher in the heterochromatin and ZNF/Repeats regions in comparison to the repressive polycomb regions.

### Large scale NOMe segmentation

GCH methylation values were first merged from both strands to calculate weighted average methylation per GpC and then smoothed using *BSmooth* (52) with `h=50000` as a smoothing window. A set of genomic regions, after visual inspection, was manually selected and labeled accordingly as S1 (not valley) or S2 (valley). These regions were used to train a random forest classifier using the average GpC methylation in 30kb tiles as variable vector and the aforementioned labels as response vector. Then the fitted model was used to predict the status of 30kb tiles across the whole genome. All predicted consecutive S2 with gap length less than 30kb were merged into one region. The final set of S2 regions were visualized in Figure 2A as valleys track. The training and prediction process were carried out using R caret package (53).

## RESULTS

### Genome wide comparison of open chromatin assay data

To monitor the comparability of the currently most widely used open chromatin assays we performed a direct data comparison between NOMe-Seq, DNase I-Seq and ATAC-Seq, outlined in Figure 1, using data generated from the same batch of HepG2 cells grown in our lab. This approach reduces experimental confounding effects such as differences in cell batches and culture condition (growth, density). Data sets were generated following existing IHEC and BLUEPRINT protocols. The HepG2 cell line was selected as it is a major cell line used in ROADMAP and ENCODE analyses further allowing us to include external datasets for subsequent analyses.

A visual inspection of all three sequencing tracks indicated a fairly good agreement of local enrichment profiles at open chromatin sites (see Figure 1B) between ATAC-and DNAse I peaks and NOMe signal enrichments. However, the NOMe-Seq coverage is much more widespread as compared to the strong local enrichments of the two nuclease-based assays. To better compare the NOMe signal distribution with ATAC/DNAse I “peak calling” we calculated the genome wide GCH methylation levels of NOMe-Seq data and related it to the FPKM values for DNase I-seq and ATAC-seq as numbers of 5’ read ends across the genome. We correlated these genome wide raw signal distributions aggregated in 500bp bins (see Figure S2). Following this approach we observe a moderate level of genome wide correlation between ATAC-seq and DNase I-seq read distributions (Spearman cor.: 0.41) while the correlation to NOMe-seq is far less only reaching 0.21 and 0.25 to ATAC and DNase I, respectively. To better understand the assay specific differences in the genome wide read distribution we started by investigating the sequence preferences of all three assays in more detail.

### Sequence, structure and DNA-methylation influence DNase I, ATAC and NOMe

In line with (12) and (39) we observed that both endonucleases DNase I and the modified Tn5 (ATAC) show a slightly biased and distinct sequence preference at their cutting sites (see Figure 3A). This preference is found across multiple samples of different cell types (see Figure S3 and S4). Also the ‘NOMe enzyme’ M.CviPI, which recognizes and methylates at the dinucleotide 5’GC3’ showed some minor sequence preferences at the flanking −2 and +2 position. However, this is only due to the (bioinformatic) exclusion of ambiguous GCG sites, which overlap with sites of endogenous CpG methylation and are therefore inconclusive. Despite the GCG effect, no other sequence biases were observed (see Figure S5).

**Figure 3.**
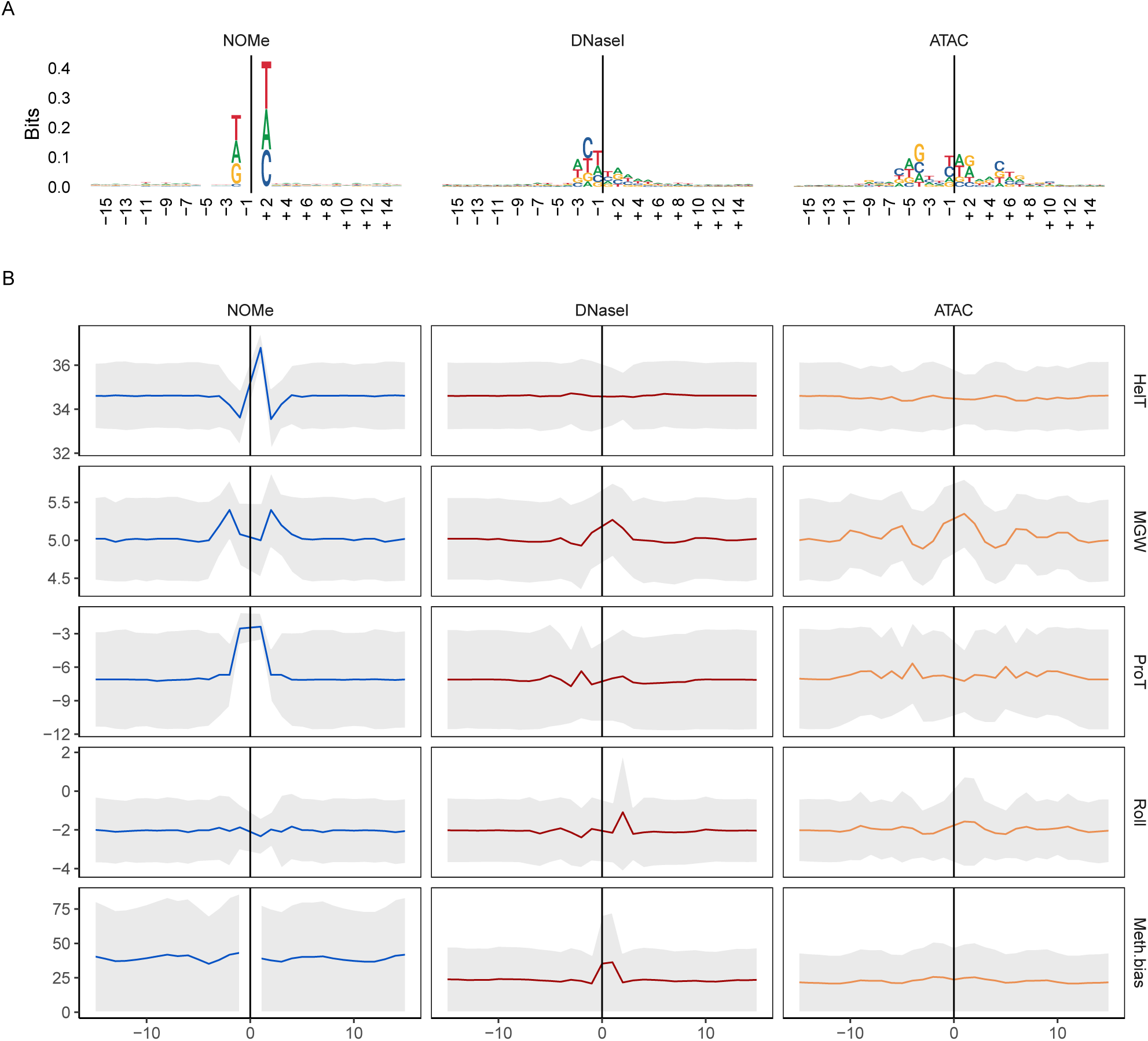
Structural and sequence preferences for NOMe (left), DNase I (middle) and ATAC (right) are shown in columns. Cutsites (ATAC,DNase I) and sites of GC methylation (NOMe) are shown as position 0 (x-axis) with a window of 31bp around it. For ATAC, a shift of 4bp upstream is introduced to conserve symmetry. A) Sequence logo for all assays, the larger a character is shown in a column the more often it occurs. B) Analysis of structural and DNA methylation features. The top four panels contain structural features; in order, helix turn (HelT), minor groove width (MGW), propeller twist (ProT) and roll. These are displayed as median with a confidence band of median absolute deviation (MAD). The bottom panel contains the average CpG methylation with a MAD confidence band.

In an earlier line of work it was shown that CpG methylation in DNase I cut sites increases cleavage efficiency by altering the DNA structure (54), which can also be predicted by looking at DNA shapes (55). To systematically compare the enzyme specific sequence preferences within each assay we predicted structural DNA shape features (see Methods) around the GpC methylation sites (NOMe) and the enzyme cutting sites (ATAC, DNase I) (Figure 3B and Figure S6) across a number of available samples and datasets (see Figure S6).

First, we investigated the structural features and observed that the sequences around the M.CviPI enzyme (NOMe) recognition site 5’GpC3’ show a pronounced signal for an increased Helix Twist (HelT) and Propeller Twist (ProT). Additionally, we find an increased Minor Groove Width (MGW) flanking the GC site. For DNase I, we observed an enlarged predicted MGW around the cut site, as reported before (54), and a slightly increased base roll. Both, M.CviPI and DNase I, act as monomers and show clear one sided effects around the recognition/cut site. The modified Tn5, on the other hand, acts as a dimer showing bidirectional oscillating changes in MGW, ProT, and Roll around the cut site. Together this analysis shows that the enzymes used in the three assays have sequence and structural preferences influencing their genome wide signal distribution.

Next, we examined if and how endogenous CpG methylation affects the three enzyme activities (see Figure 3B, bottom panel). DNase I indeed showed a slight but focused increase of CpG methylation around cutting sites - an observation confirming findings made by (54), while both M.CviPI and the modified Tn5 showed no position-specific effect of CpG methylation along the cutting site. However, we noticed that the average level of 5mC around DNase I and modified Tn5 (ATAC) cutting sites is strikingly lower as compared to M.CviPI (NOMe). This is a reflection of a much broader almost uniform (see Figure 1B) genome-wide coverage of NOMe reads.

### Shared and specific features of nucleosome depleted regions recognized by NOMe, DNase I and ATAC assays

Next we focused our attention on comparing the performance of all three methods in open chromatin sites or Nucleosome Depleted Regions (NDRs). Such regions are widely used for functional genome annotations and interpretations. For NOMe we calculated NDRs with our own HMM based approach called gNOMeHMM (see Methods,(56)) which provides a robust genome wide NDR annotation (see Methods).

NDR detection for DNase I-seq and ATAC-seq data was performed with MACS2 (see Methods). Overall, we detected very similar numbers of NDRs for NOMe-seq (65,683), DNase I-seq (62,365) and ATAC-seq (67,675). Note that for all three assays libraries were sequenced at a depth to obtain sufficient coverage (NOMe = 10 x genome wide coverage of GpCs), DNAse I 180 million reads, ATAC 60 million reads, see Methods). The total number of NDRs recognized by at least one of the three assays sums up to 105,081 of which most were found in intergenic (*∼*55%) or intronic (*∼*25%) regions. In total, 24% of NDRs were shared by all methods and additional 27% were supported by two methods (see Figure 4A).

**Figure 4.**
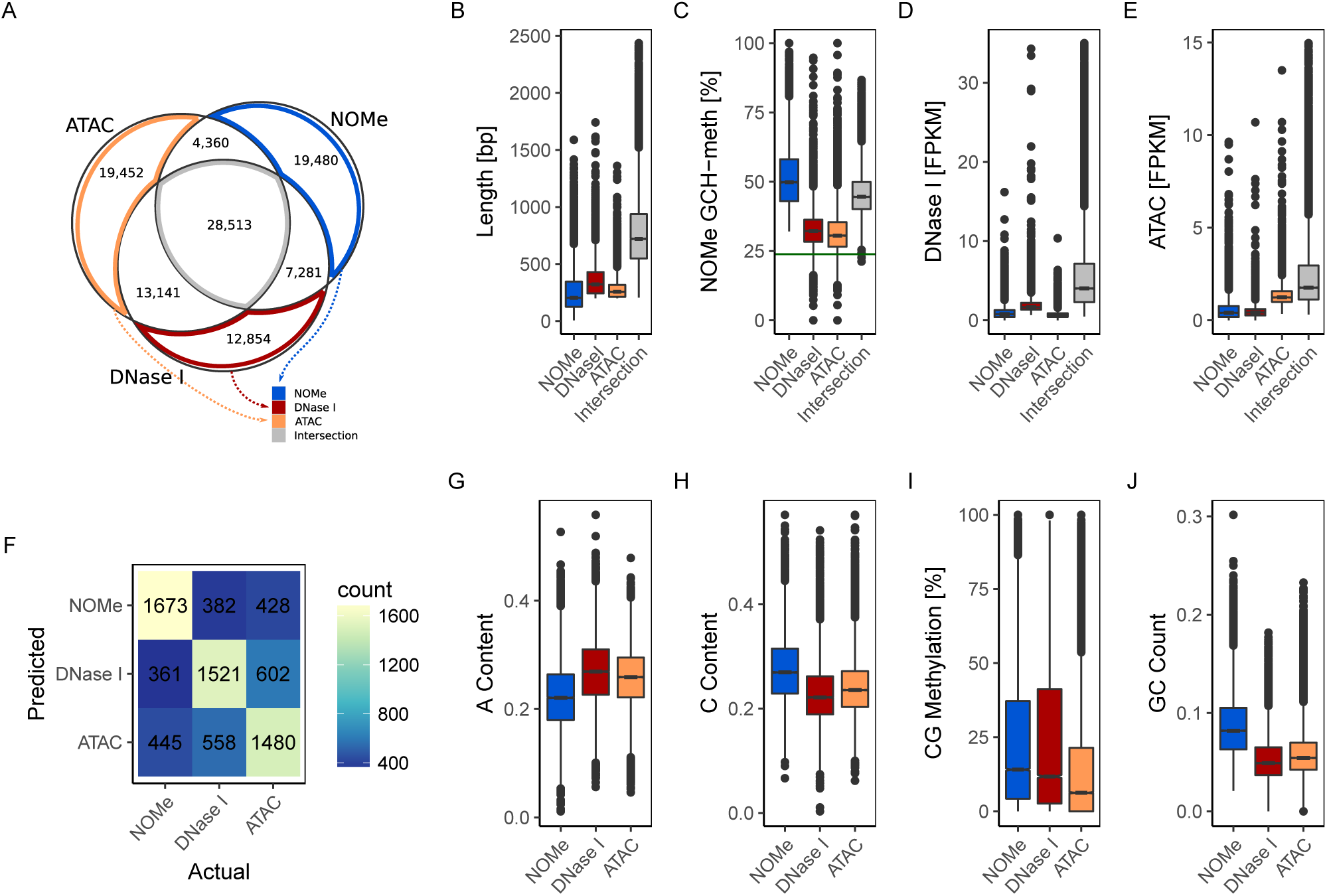
Comparison of accessibility measured with DNase I, ATAC and NOMe. A) Comparison of NDR calls by all three methods, see text. B) length distribution of NDRs. C) NOMe signal at NDRs. D) DNase I signal at NDRs. E) ATAC signal at NDRs. F) Results of classification of assay-unique NDRs. The confusion matrix shows actual (column) against predicted (row) class labels. G-J) box plots of sequence characteristics selected by the classifier to have power in separation.

19,480 unique NDRs were predicted for NOMe, 12,854 for DNase I and 19,452 for ATAC. Most shared NDRs were found between ATAC and DNAse I data, underpinning the commonalities in the enzymatic (cutting) reaction of DNase I and ATAC and also probably the commonalities in data processing and peak calling.

Common NDRs, i.e. NDRs shared by all three assays, tend to be longer, largely overlapping with previously known NDRs (see Figure 4B and S7B). They, in general, exhibit a strong above-average signal in all assays. NOMe-alone NDRs are an exception to this rule with slightly stronger signal as compared to common NDRs. Common NDRs are most strongly enriched for TSS, followed by CpG islands (CGIs) and LaminB1 sites and are heavily enriched for ENCODE mapped transcription factor binding sites (TFBSs) (see Figure S7B). Assay-unique NDRs (ATAC-alone, DNase I-alone, NOMe-alone) have a relatively strong signal in only one of the data sets and the signal intensity drops or disappears in the other two assays (See Figure 4C-E). LOLA annotation of such assay-unique NDRs shows a specific enrichment of NOMe and ATAC unique NDRs for CTCF, Rad21 and SMC3 binding sites. This enrichment is even more pronounced in NOMe-unique NDRs. Rad21 and SMC3 are parts of the cohesin protein complex interacting with CTCF (57). Their co-localization suggests a widespread distribution of dynamic topological substructures (58) preferentially detected as NOMe and ATAC NDRs, respectively. Unique DNase I NDRs on the other hand are enriched for binding sites of the hepatocyte nuclear factors like FOXA1, FOXA2 and HNF4G indicating a higher sensitivity of DNAse I to recognize networks of liver-specific NDRs.

### Sequence features and methylation enrichment around NDRs

To better characterize features separating the three sets of unique NDRs we applied a logistic regression classifier. The feature set included DNA methylation and sequence associated measures (see Methods). The accuracy of the classifier was 0.55 on a hold out test data set. Note that, as this is a three-class classification problem, a random classifier would achieve an accuracy of 0.33. When we additionally included the counts of 5-mers in the regions, the accuracy increased to 0.63. The most important features were A-content, C-content, GC count, and CG methylation (see Figure 4G-J). The former three separate NOMe from the enrichment assays, while CG methylation helps distinguish ATAC from NOMe and DNase I (list of top features shown in Table S2). Additionally, some 5-mers were useful in separating the classes. For example, for classifying NOMe unique NDRs the 5-mer `CGCGC` was depleted, representing the GCG effect, while 5-mers enriched in DNase I unique NDRs resembled the observed DNase I sequence bias. Overall, the classifier was better at separating NOMe from ATAC and DNase I, and the most miss-classifications occurred between the ATAC and DNase I classes (see Figure 4F). The link between CG methylation and DNA accessibility is known (59). While only 10% of the Intersection NDRs have a CG methylation above 30%, there are 20% unique ATAC NDRs and about 30% unique NOMe and DNase I NDRs that fulfill this criteria (see Figure S8A).

In addition, we looked to sequencing coverage and copy numbers (e.g. low complexity repeats) on NDR calling. Most NDRs are gathered around the median 16x read coverage, but there is a long tail towards high coverage. About 2-3% of unique NOMe NDRs had a coverage in the top 5%, while NDRs only detected with ATAC and/or DNase I had 8-9% (see Figure S8B). This is concordant with a higher fraction of NDRs with at least 50% overlap to repeat masked regions among unique ATAC (38%) and DNaseI (34%) NDRs as compared to unique NOMe NDRs (26%) and the intersection (15%) (see Figure S8C). A feature singling out unique NOMe NDRs is the distance to the next NDR in the union set; 14% of unique NOMe NDRs are within 200bp, while the next highest value is about 8% for other subsets containing NOMe NDRs in Figure 4A (see Figure S8D).

### Nucleosome phasing around NDRs

We analyzed the extent of nucleosome phasing around NDRs (see Supplementary text). While DNase I strongly uncovers the first nucleosomes 5’ and 3’ around the NDR, NOMe shows a nice “oscillating” pattern of up to five nucleosomes around many NDRs (see Figure S9). Also ATAC-seq has been shown to allow the detection of phased nucleosomes provided sufficient sequencing depth (12). In our setting ATAC-Seq with 60 million reads nucleosome phasing around NDRs was only visible when fixing the summit of NDRs called by ATAC or at borders of NDRs called by NOMe (see Figure S9). Considering that the actual sequencing coverage of NOMe-, DNAse I-and ATAC-seq around NDRs is comparable, the detection of phased nucleosome patterns is much more pronounced for NOMe-Seq. Reasons for this might be read information density and the genome wide coverage of NOMe-seq (see Supplementary text.

### NDRs can be grouped based on assays, size and nucleosome phasing

Next we performed K-means clustering of all NDR’s in a 2kb window centered around the NDR peak (see Figure 5A). We included all regions providing a NDR signal in at least one of the assays. The compilation of all NDRs illustrates that NDR size, signal strength and nucleosome phasing are prominent characteristics of both assay specific and assay independent patterns contributing to the formation of 15 NDR clusters. NDR clusters 3, 12, 13, and 15 are strongly enriched for TSS associated regions. They are marked by strong NDR signals mostly shared across all assays. These NDRs are characterized by an enrichment for general TFBS such as GABP, TAF1, and TBP (see Figure 5B). Clusters 10 and, to a lesser degree, 11 show a similar enrichment for TFBS but lack the strong TSS enrichment. Clusters 3 and 10 show an enrichment for TBP and bidirectional promoter activity. Cluster 1 and 6 have a moderate enrichment for active enhancers of class 1, also showing an enrichment for FOXA1, FOXA2 and to some degree HNF4G, i.e. a set of liver-specific transcription factors. NDR clusters 4 and 7 are most prominently called by NOMe-seq data. They were mildly (4) or strongly (7) enriched for CTCF TFBSs. Both clusters showed a regular distribution of open and closed chromatin signals indicating strong and extended nucleosome phasing around CTCF sites as reported by (22).

**Figure 5.**
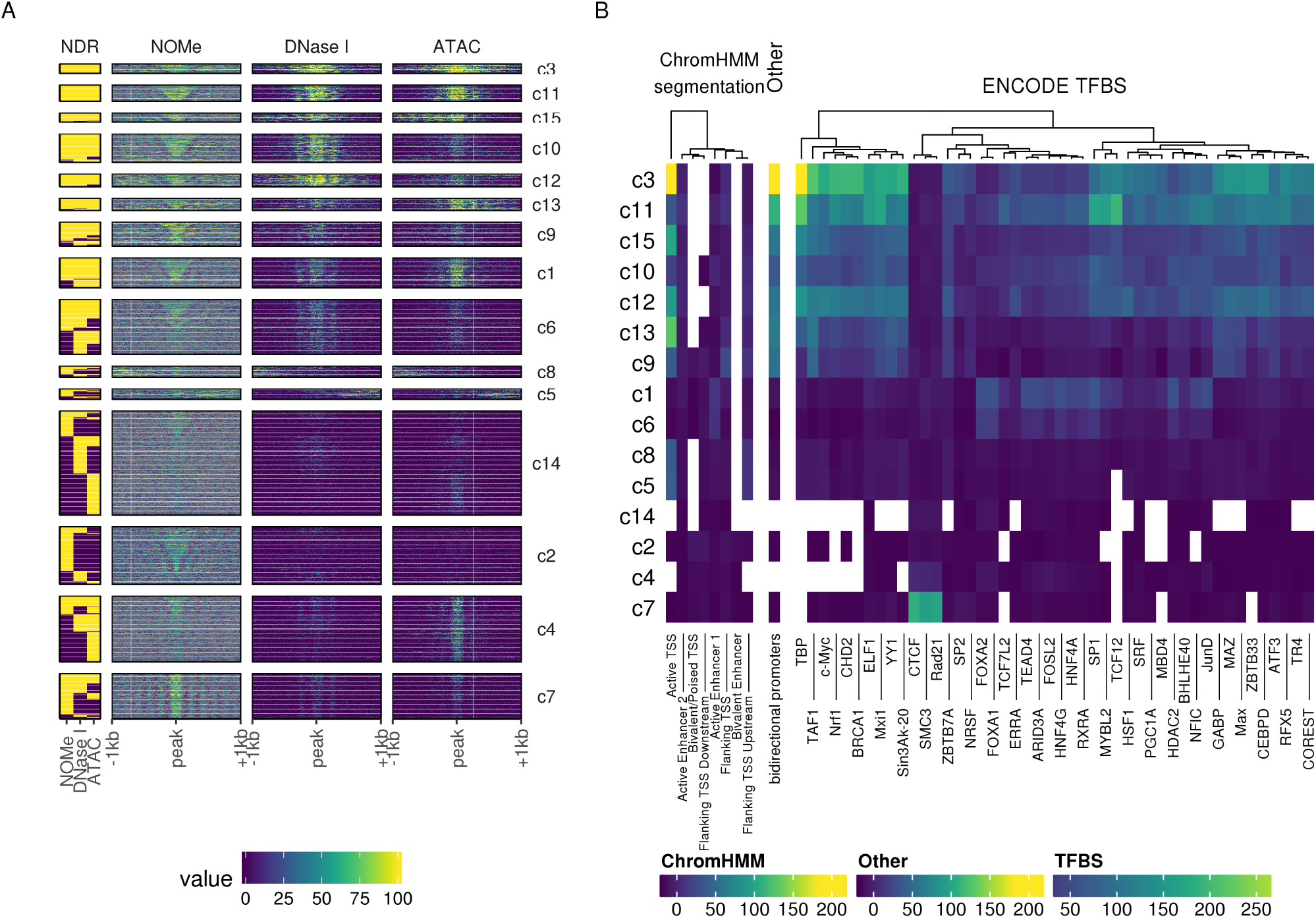
Annotated clusters of NDRs with similar signal profiles. A) NDRs signal profiles clustered into 15 clusters with K-means clustering. The left-most column shows the presence (yellow) of a called NDR for each method, respectively. For each assay, 2kb centered on the NDR was split into 10bp windows over which the signal was aggregated. With NOMe the raw methylation is displayed, while the DNase I and ATAC are represented by normalized log2-scaled read counts. B) Annotation by LOLA of each cluster. The coloration of each tile corresponds to the log odds ratio of the enrichment test. These tests were only conducted against HepG2 tracks.

NDR clusters 4 and 7 also hold among the most narrow NDRs, which is in agreement with observations that nucleosome distance around NDRs with CTCF binding sites is smaller compared to, for instance, FOXA1 sites (see Figure 6A). Cluster 14 aggregates low signal NDRs, characterized by a lack of TFBS enrichment but an increased fraction of highly methylated and repeat masked loci. Together the clustering shows that parameters such as size, signal strength and nucleosome phasing are linked to functionally distinct NDR classes across the genome as also reported by (60). Particularly NOMe-seq and ATAC-seq show an enhanced sensitivity for detecting distinct groups of NDRs outside of annotated enhancers and TSS.

**Figure 6.**
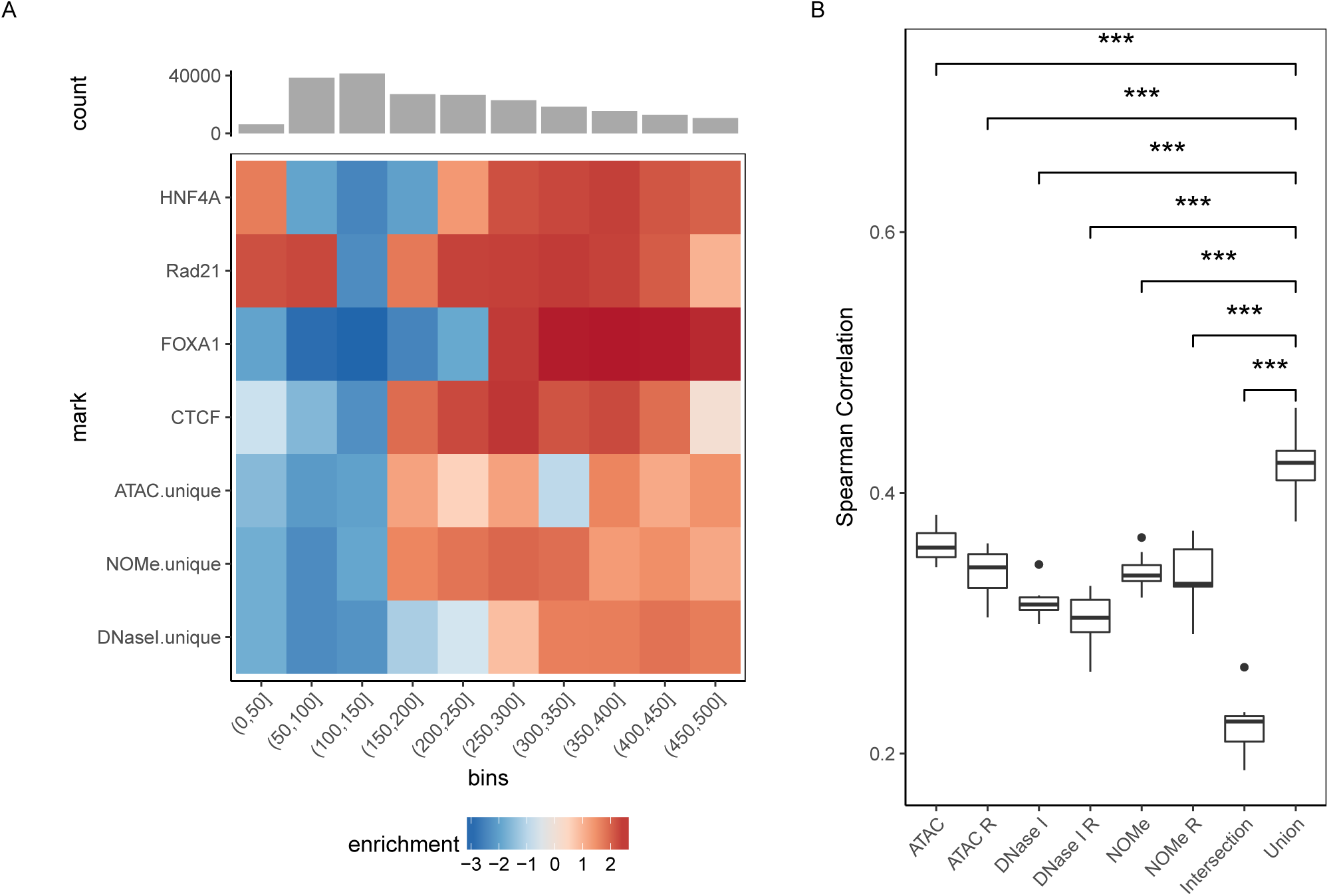
Functional analysis of shared and assay-unique NDRs. A) Analysis of nucleosome distances overlapping NDRs unqiue to any of the three assays as well as ChIP-seq peaks for the TF CTCF, RAD21, HNF4A, FOXA1 (rows). Intensity in the heatmap encodes enrichment compared to the background distribution in regions of the same length (column). B) Comparison of gene expression prediction using TEPIC when using different NDR sets as basis for feature calculation (x-axis). The assay-specific NDR sets for ATAC, NOMe and DNase I are compared to the union or intersection NDR sets using all assays. In addition, each assay-specific NDR set was increased by adding random regions of the same size, as compared to the union set, denoted as ATAC R, NOMe R and DNaseI R, respectively, to assess the effect on prediction performance with increasing set sizes. Gene expression prediction performance is shown as a boxplot of Spearman correlation values (y-axis) between true and predicted expression values as part of 10-fold cross validation on genes not used for learning. *** t-test p *<* 0.01.

### NDRs can be used to predict expression level of nearby genes

We recently developed *TEPIC* as an open chromatin based prediction model for gene expression using predicted TFBSs (45). With this approach we compared the prediction performance for NDRs individually called by ATAC, DNAse I, and NOMe, as well as the intersection and the union of those (see Figure 6B). For each NDR subset, we computed TF binding predictions for 726 TFs and used these predictions in a gene-centric way as features to predict gene expression (see Methods). The worst performance was obtained using the NDRs obtained by all three assays (intersection including 28,513 NDRs), which led to smaller performance Spearman correlation values although many of these NDRs share a strong signal in all assays. Note that these shared NDRs mostly cover TSS and active enhancers. For NDRs specific (not unique) to each assay, we observed that the results are comparable between assays with a slightly better performance of ATAC over NOMe and DNase I.

The best model performance, contested by no other set, was achieved by combining all three NDR sets (union). To assure that this is not simply due to the increased number of NDRs, we extended each assay specific NDR set with randomly generated peaks to match the size of the union set. These models performed constantly worse than the actual assay specific sets and thus also worse than the union set. Put together this suggests that each assay fails to describe a certain part of the accessible chromatin landscape and thus, while the combination of assay specific NDRs allows to more comprehensively model the regulatory influence on gene expression.

### Confirmation of assay specific NDRs

To verify the assignment of common and unique NOMe, DNase I and ATAC NDRs by an independent assay we selected 17 NDR regions showing distinct recognition patterns across the three assays (Figure S10A). We analyzed these regions by targeted deep amplicon bisulfite sequencing following a NOMe treatment in HepG2-cells (see Figure S10B for the experimental data of four examples). For NOMe-unqiue NDRs we observed a median GCH-methylation level of 35-70% (with the exception of the Fam35DP amplicon), while regions not called from NOMe-Seq data (no NDR in any of the three methods or NDRs unique to DNase I and/or ATAC) show a reduced median GCH methylation of 9-26% (Figure S10). Overall this confirms that NDRs are reliably called by gNOMeHMM showing a clear and specific GCH enrichment profile above background (except for FAM35DP with median GCH-methylation of 17%). Still, we also detect intriguing patterns of GCH-methylation in regions not called by gNOMeHMM, but only by the other methods. One such example is the DNase I-and ATAC-unique NDR NCK2 (Figure S10B) and the other a non-NDR region within the UGGT1 (data not shown). Both show a clear GCH-methylation pattern in a subset of sequences indicating that a proportion of cells fulfill the criteria of a NOMe-specific NDR. This finding illustrates that some (partially) open chromatin regions might be missed by gNOMeHMM calling either due to low GCH-density or by low GCH-methylation levels indistinguishable from the neighboring “background signals”.

### Genome wide information called by NOMe

In comparison to ATAC and DNase I, NOMe-Seq data show an almost complete coverage across the genome. Outside of NDR peak regions this has mostly been treated as background noise. Here we had a deeper look into chromatin associated features linked to this genome wide “background”.

#### Nucleosome phasing around Intron-exon junctions

We first analyzed the nucleosome phasing in a 2 KB window around intron-exon junctions for all genes. In short we merged all NOMe data around intron-exon boundaries, excluding the first and the last two exons. To our surprise we detected a very pronounced phasing of up to 1Kb around the exon/intron junctions in genes with low expression when we applied K-means clustering (n=5). This could not be observed for neither the unexpressed genes nor the moderate to highly expressed genes (see Figure S11). Since expression is known to be related to histone elongation marks (61) we therefore analyzed the distribution of histone marks in the clusters in relation to phasing. Analogous to (62) we observe some reciprocal distribution of histone modifications at introns and exons (e.g. higher H3K36me3 in moderate to highly expressed exons) but no obvious direct link to the clusters showing phased or non-phased nucleosomes.

#### Genome wide distribution of NOMe signals in relation to chromatin states

While NOMe-Seq has originally been used to assess chromatin accessibility, additional information content in the genome wide GpC methylation has largely been ignored. Visual inspection of GpC methylation in HepG2 cells indicated a minor variation of “background” GpC signal across large domains (see Figure 2A-B), here called valleys. We therefore systematically analyzed the GpC methylation across 18 chromatin states for a series of cell types (HepG2, monocyte, machrophage, CD4 T central memory, CD4 T effector memory and CD4 T naive). We noticed that while the general level of genome wide GpC methylation changes with samples, the relative distribution over chromatin states remains constant (see Figure S12). In fact, the lowest GpC signal was always in the *Heterochromatin* and *ZNF genes & repeats* states of the Roadmap 18-state ChromHMM segmentation.

In HepG2, we identified valleys using a classification model (see Methods). Surprisingly, the valleys are largely concurrent with partially methylated domains (PMDs), as defined by (49), and B-compartments retrieved from the Hi-C data of HepG2 (see Figure S13). Moreover, the strongest segmental decrease of GpC methylation is co-localized with the most compacted heterochromatic domains, as measured by DLR score (Distal-to-local Ratio) based on Hi-C data (see Methods) (see Figure 2B-C).

#### Accurate genome wide CpG methylation calling

NOMe-Seq has the great advantage to not only determine the genome wide distribution of closed and open chromatin but to also provide detailed information of the genome wide “endogenous” CpG methylation status. To monitor this in detail we performed a direct comparison between NOMe-seq data and an independent WGBS-dataset generated from the same HepG2 cell batch. NOMe-seq called “endogenous” CG methylation correlated excellent with the conventional WGBS data (Pearson correlation 0.95). This excellent correlation was observed on the level of single-CpG methylation as well as aggregated methylation levels at promoters, CpG islands and genome-wide tiled regions (see Figure S14B-C). We noticed, however, that when confining our analysis to aggregated methylation levels in the regions identified as open chromatin by NOMe-seq (NDRs see below), correlation values slightly decreased as compared to genome-wide analysis, potentially due to the general depletion of CpG methylation in open chromatin (see Figure S14B,D). We also observed a tendency of a decreased NOMe-seq read coverage at regions exhibiting the highest degree of differential methylation (see Figure S14E). Interestingly, the highest ranking differentially methylated regions appear to have a higher CpG dinucleotide content whereas the GpC dinucleotide content is more uniformly distributed across all open chromatin regions (see Figure S14E). We conclude that NOMe-seq data provide an overall excellent resource to determine endogenous CpG methylation largely independent from the GpC measured chromatin accessibility (see Figure S14).

In summary, NOMe-Seq provides not only openness/closeness information at NDRs such as TSS, enhancers or other short regulatory regions, but also includes additional information about the genome wide chromatin compaction. Such additional features detectable in NOMe “bulk” sequencing data sets may be useful for the interpretation of genome wide single cell NOMe-Seq data.

## DISCUSSION

There are many distinct technical approaches to map open chromatin such as MNase-seq (63, 64), FAIRE-seq (65), SONO-seq (66) but also using ChIP-seq data as reference (e.g. NucHunter (42)). Here, we focused on the three most commonly used methods ATAC, DNase I and NOMe, and thoroughly compared them in respect to their commonalities and differences as well as individual strengths and weaknesses.

### Experimental differences and open chromatin calling

While all three methods require proper isolation of nuclei, the experimental challenges for the three assays differs. ATAC has the fewest working steps and nicely links labeling of accessible regions and NGS library preparation by the transposase-assisted incorporation of NGS adaptors into open chromatin. For DNase I it is important to avoid/minimize loss during the isolation of the small double-cut DNA fragments and the NGS library preparation. The experimental challenges for NOMe are comparable to DNase I although it is not an enrichment assay, but rather an extension of the commonly used bisulfite sequencing. This makes it possible to read out native CpG-methylation in addition to the chromosome accessibility and in scenarios where information from both epigenetic layers is desired, NOMe kills two birds with one stone. Since each NOMe sequencing read potentially reports on several linked events from a single cell, it provides an excellent opportunity to detect sub-population effects. With DNase I and ATAC, these events are commonly used to detect TFBS by genomic footprinting (67, 68). Although, the coverage is relatively limited for genome-wide NOMe, such a procedure can be bolstered by multiple measurements from single cells as seen for CTCF (22). Our analysis also shows that irrespective of how NDRs are called NOMe allows (for most) a straight forward and very sensitive and quantitative way to deeply analysis the local chromatin accessibility via ultra deep sequencing of bisulfite amplicons.

### Assay specific variation in NDR detection

A qualitative and quantitative interpretation of nucleosome depleted regions (NDRs) also called “open chromatin sites” is the most valuable information deduced from chromatin accessibiltity data. Not surprisingly NDRs detected by all methods tested display strong signals. They are strongly enriched for TFBS and mainly cover active TSSs, common enhancers and other shared accessible/open loci (see Figure 5 and Figure S7).

However, this core of strong and well-characterized NDRs is apparently less informative to predict gene expression, hinting towards the importance of weaker and more difficult to detect cell type specific NDRs outside of the intersection (see Figure 6B). NDRs singularly or dually detected by the individual assays outperformed the intersection in expression prediction. Indeed the best predictions were made with the union of all NDRs indicating that each set of unique NDRs carries important extra information not covered by the other assays.

To understand the differences in detection between the assays we investigated unique NDRs, i.e. NDRs called by one assay only, in more detail. Although these NDRs were not called by one of the other two assays (note that we applied conventional sequencing depth for all assays), a deep NOMe-Seq analysis suggests that seemingly unique NDRs can in principle also be detected by other assays (at least in one direction). Moreover enrichment analysis of known NDRs (see Figure S7), suggests that most of these (unique) NDRs are trustworthy. However overall the unique sets of NDRs fall into two groups: DNase I NDRs are higher enriched for cell type (i.e. liver specific) enhancers overlapping with FOXA1, FOXA2 and HNF4G binding sites, while NOMe and ATAC NDRs often demarcate insulator regions associated with CTCF, Rad21 and SMC3 binding sites. Together these findings argue for a more careful assay specific and context dependent interpretation of open chromatin maps generated by only one assay and their limited use for trans-assay comparative analyses.

A potential reason for a preferential calling of insulator regions by ATAC and NOMe (see Figure 5) in our setting appears to be the variation in fragment length distribution between assays (see Figure 6A). DNase I-Seq libraries are known to be dependent on enzyme concentration and insert size selection for library construction (10). NOMe and ATAC assays are used as “endpoint” reactions and libraries are generated without size preselection. In our comparison they indeed cover a relatively smaller size range (see Figure 6A) such that the enrichment of relatively short insulator NRDs may be simply due to preparative differences.

Moreover, each assay comes with a set of inherent limitations affecting NDR detection. NOMe for example is only able to measure open chromatin in regions containing a sufficient 5’GpC3’ sequence context. Simultaneously, as shown by (12), (39) and confirmed here (see Figure 3, S3 and S4), the enyzmes used for DNase I and ATAC assays come with a slight sequence preference. Here, we show that also the tertiary structure of the DNA has an apparent influence on enzyme activity (see Figure 3). Moreover there is an influence of endogenous 5’CpG3’ methylation surrounding the enzyme activity site. The increased methylation level at DNase I cut sites could be a sign of how at least one of these reside outside the NDR. In both NOMe and, especially, ATAC we observe a weak periodic pattern in the methylation bias that coincides with the minor groove width (see Figure S6). It could be speculated whether this is due to how DNA is wrapped around the nucleosome as described by (59).

With NOMe, we observed more NDRs close to other stronger NDRs (see Figure 5, cluster 5 and 8). These NDRs are probably not unique NDRs, but rather a result of phased nucleosomes and hence a sign of the strength and homogeneity of the stronger NDR. Our suggestion is to handle flanking NDRs with care and assign extra weight to the focal NDR at such a loci. For the enrichment assays — DNase I and ATAC — a potential sign of false positives would be high-ploidy or repeated regions artificially amplifying coverage and hence generating peaks. A common strategy to avoid this and other biases is the sequencing of a non-enriched library (cf. input sequencing for ChIP-seq). The most important factors, when applying machine learning to the classification problem of whether a NDR was unique to either assay, was A-, C-content, GC count and native CpG methylation.

### Unique epigenomic information provided by NOMe

DNase I and ATAC-Seq data are the most cost efficient and widely used techniques almost exclusively used for the local detection of open chromatin sites. In comparison to NOMe they only require a moderate level of sequencing depth and ongoing improvements in protocols are reducing that depth even further. However, what is often neglected is that the higher sequencing expenses for NOMe come with much deeper information content. First and most importantly in comparison to all other enrichment technologies (including MNase, FAIR and others), NOMe-Seq provides a single chromosome readout of multiple linked measurements allowing a direct localization and quantification of epigenetic changes. Recent developments in NOMe-Seq by (69) strongly argue for the use of NOMe for single molecule footprinting. Secondly, NOMe not only provides information of open chromatin sites (NDRs), it also allows to call endogenous DNA-methylation at the same time. These features have been appreciated by several authors making NOMe-Seq a prime method for deep single cell epigenomics (4, 5). However, so far most analyses concentrated on NDR detection and endogenous methylation calling, neglecting other genome wide informations we started to decipher in our analyses. Our extended study shows that GpC methylation profiles provide a measure for the extend and distribution of heterochromatin compartments and also the local nucleosome phasing outside of NDRs e.g. around exon/intron junctions of genes across the genome. Surprisingly, this local phasing had some relation to gene expression strength. Additional work will be needed to understand the functional link between regular nucleosome spacing and low gene expression. We believe that such additional features are important add-ons that currently can comprehensively only be achieved by NOMe-Seq.

In conclusion, our controlled side by side comparative approach of open chromatin assays revealed that all three most commonly used assays allow to call the most prominent NDRs covering a large fraction of the (mostly functionally annotated) highly accessible regions. However, we also find that many NDRs are less likely to be detected/called by individual assays and that these additional assay unique NDRs are extremely important to predict (and hence explain) the expression of genes. Our findings suggest that single assay approaches to detect open chromatin are less comprehensive than anticipated and incomplete for the calling of regulatory open chromatin information (at least under standard settings). Finally we show that assays such as NOMe-Seq that are more sequence consuming, but cover the genome in a comprehensive manner provide information on a series of useful additional epigenomic features important for functional interpretation.

## Supporting information

Table summarizing data sets used

## DATA AVAILABILITY

All data sets used in this work are listed in supplementary file.

## SUPPLEMENTARY DATA

dataSources.xlxs: Spreadsheet containing sources for data used in this work

## AUTHOR CONTRIBUTIONS

KN, FS, GG, FM, PE, MS and JW contributed to Project conception. KN, FS, AS, FM, PE, IGC, NP and MS contributed to the bioinformatic analysis. KN, FS, NG, AS, GG, KK, PE and JW contributed to the experimental analysis. NG, GG and KK contributed to the primary data generation. KN, FS, NG, AS, GG and FM generated the figures. KN and JW drafted the article with the help of FS, NG, AS, GG, FM, NP and MS.

## SUPPLEMENTARY TEXT

### NOMe NDR prediction based on hidden Markov models

As NOMe is a bisulfite based assay the signal is qualitatively different compared to ATAC and DNase I Nucleosome

Depleted Regions (NDRs) we cannot utilize conventional peak callers. Previous analysis of NOMe data was based on the window-based method findNDRs (22), that computes enrichment of methylated reads compared to the genome average in a window of fixed size with a Chi-square statistic. findNDRs has the disadvantages that the minimal size of a detectable NDR is set by the user and enriched windows are merged using a heuristic average of p-values. Finally, significant NDRs are estimated based on a hard cutoff on the p-value derived from a small set of NOMe datasets without proper control for multiple testing. Here, we suggest the first method to call NDRs from NOMe data based on a hidden Markov model, separating putative NDR regions from background noise. The method, called gNOMeHMM, works with methylation levels of individual GCs in GCH context and is therefore able to find NDRs of arbitrary size. In a second step, significance of enrichment is contrasted to a background model based on the local surroundings rather than a fixed global average. Another advantage is that each NDR is associated to a robust p-value corrected using an empirical false discovery rate (FDR), see Methods. One pitfall with p-values generated by Fisher’s exact test or the Chi-square statistic is that they are heavily coverage dependent. While high coverage normally is equal to greater statistical power, it can also be artificially induced by collapsing reads originating from copy number variations or other large scale genomic rearrangements to a single locus. In this case, a fixed p-value cutoff enriches for NDRs in regions with higher coverage. Our method approaches this by stratifying NDR regions by coverage through automatic clustering, see Methods. As a measure of performance we compared gNOMeHMM and findNDRs to the union set of NDRs from ATAC and DNase I data. We used our own HepG2 sample with NOMe, DNase I and ATAC experiments conducted on the same batch of a cell line in the same lab. gNOMeHMM detected 65,683 NDRs at a FDR cutoff at 0.01 and a F-score of 0.42 (see Figure S15). Relaxing the FDR threshold to 0.05 resulted in 128,837 NDRs with a conserved F-score at 0.42. The optimal F-score is found in a local maximum between the two conventionally used cutoffs; at FDR¡=0.028 with 91,202 NDRs and a F-score of 0.43. In order to avoid false positive we decided to put our continued focus on the stricter set selected with a FDR cutoff at 0.01.

### Considerations when observing phased nucleosomes

For NOMe and ATAC it is possible to observe the pattern of phased nucleosomes by characterization of regions flanking the NDRs (12, 70). A big difference between NOMe and the enrichment based assays, ATAC and DNase I, is the information density per sequenced fragment. The ATAC and DNase I assays rely on preferential cleavage of accessible DNA, and hence, each fragment carries information about two such events. With fragments long enough to cover multiple nucleosomes it is possible to infer additional cut sites within each long fragment (12). This can add one or two inferred data points for a small subset of fragments. NOMe on the other hand is dependent on the GCH density. Given our experiment in HepG2 and sequencing of 2×100bp we have around eight data points per fragment.

In order to observe conserved nucleosome phasing between loci, the choice of reference point is of utmost importance. Two intuitive alternatives are the summit of the NDR peak or the boundaries of the NDR. Here, we defined the NDR summit as the center of a NOMe interval or as the MACS2 defined summit for ATAC and DNase I. While the summit is sensitive to the actual length of the NDR, fixating on the interval boundaries requires accurate prediction of these. Comparison showed that gNOMeHMM predicts borders well, that the summit is the better reference point with MACS2 and how a poor reference point can obscure the phasing in an aggregated signal (see Figure S9).

## SUPPLEMENTARY FIGURES

**Figure S1.**
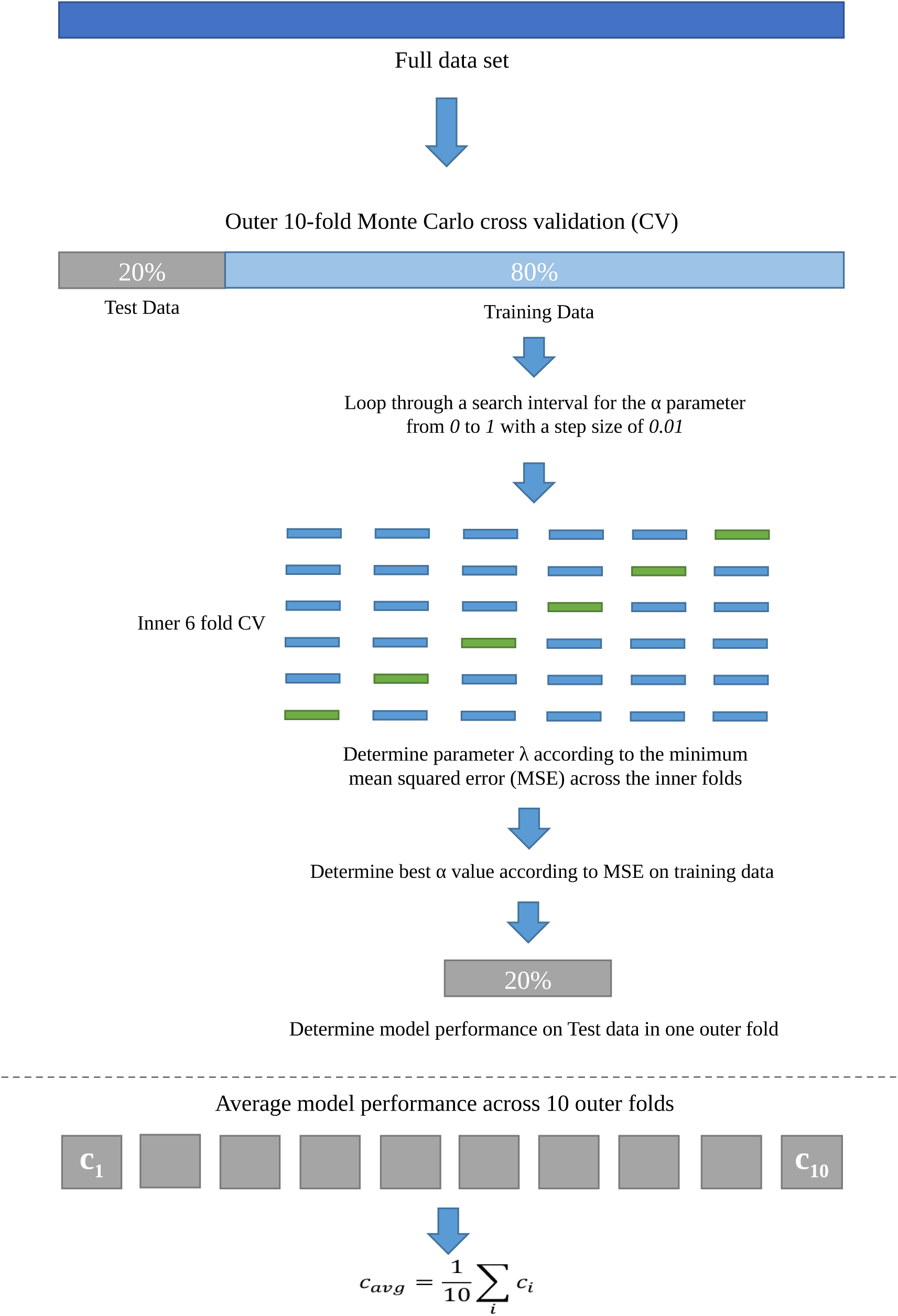
Summary of machine learning procedure.

**Figure S2.**
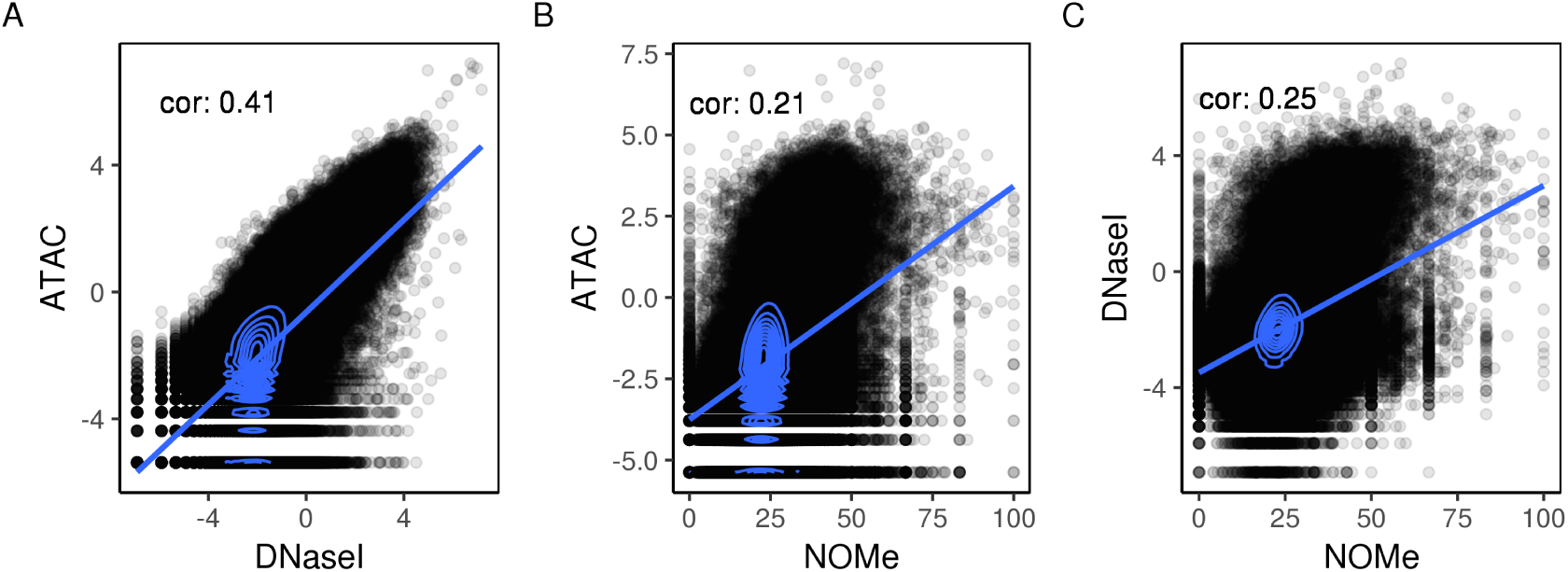
Pairwise comparison of binned signals for A) DNase I and ATAC, B) ATAC and NOMe, C) DNase I and NOMe.

**Figure S3.**
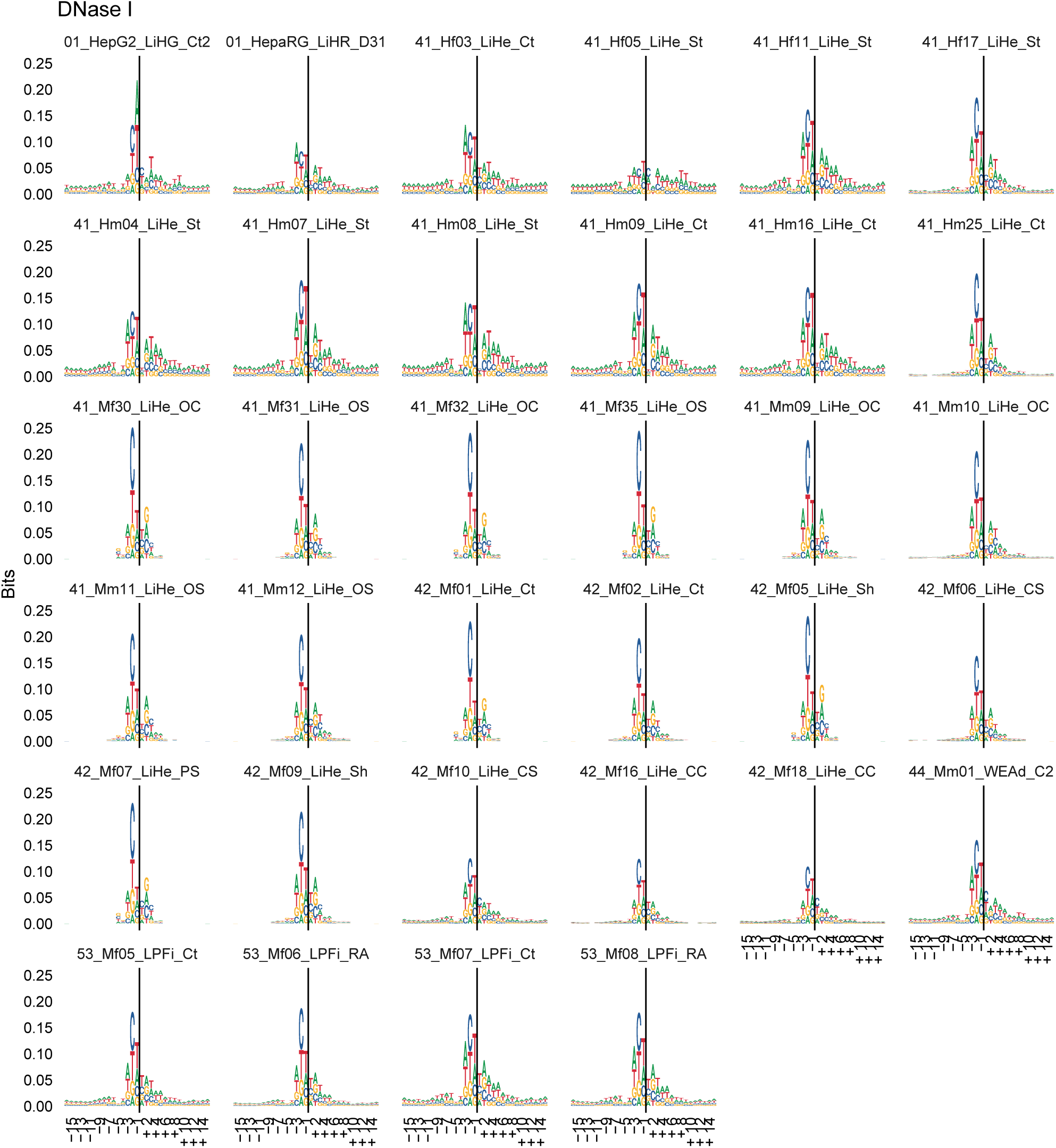
Sequence logos for a set of DEEP DNase I samples similar to Figure 3A.

**Figure S4.**
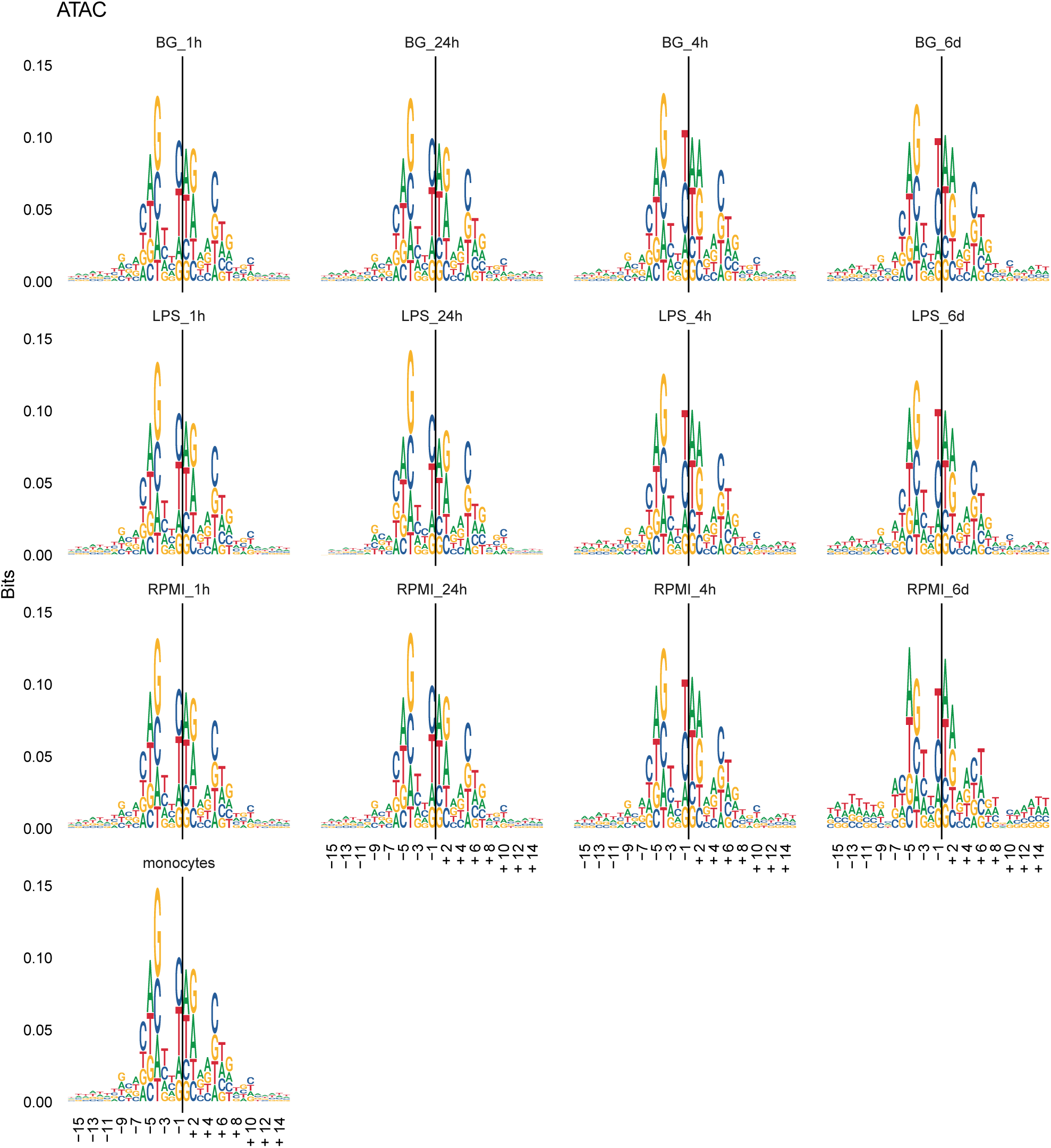
Sequence logos for a set of Blueprint ATAC samples similar to Figure 3A.

**Figure S5.**
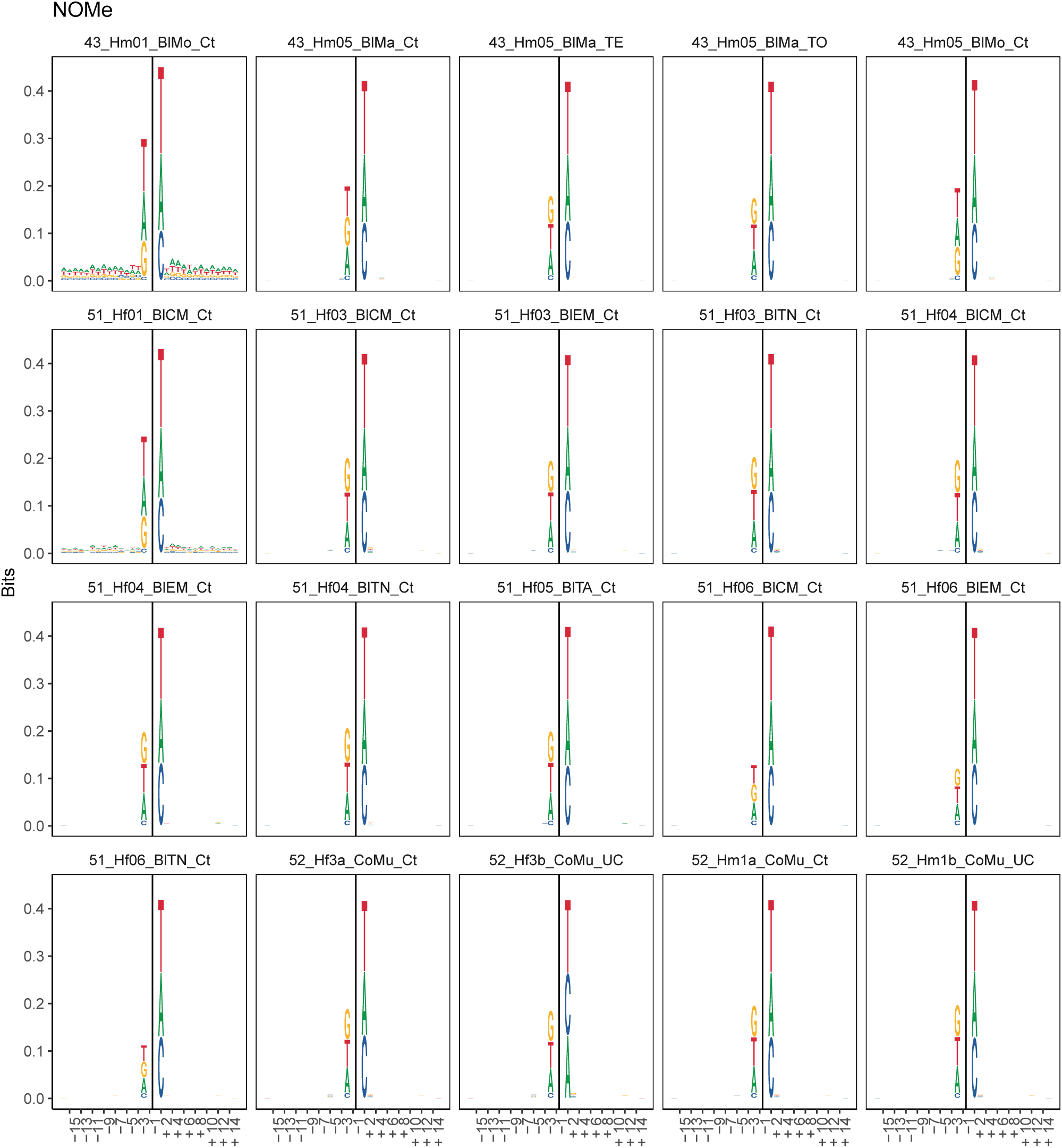
Sequence logos for a set of DEEP NOMe samples similar to Figure 3A.

**Figure S6.**
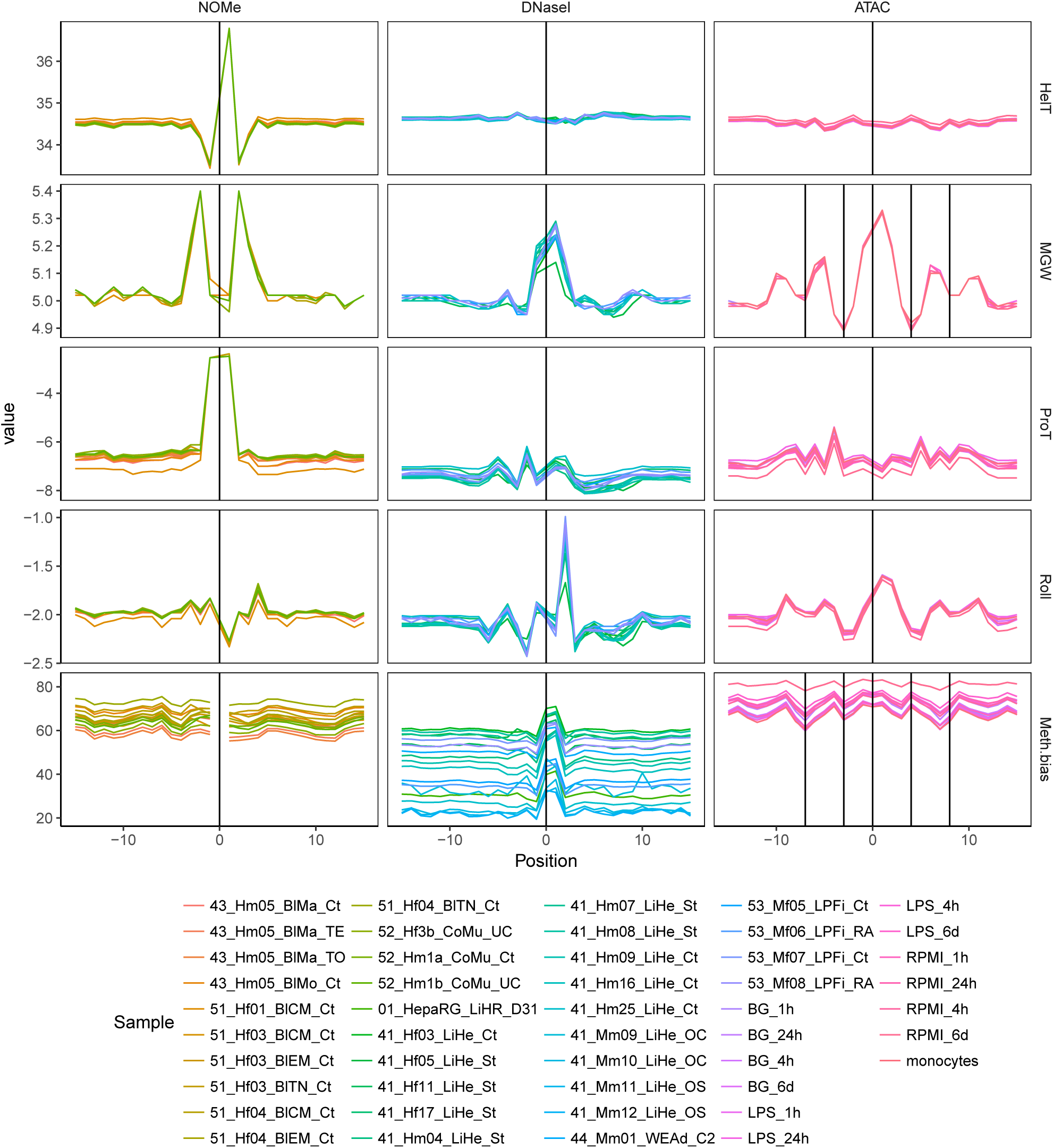
DNA structure and CpG methylation bias around enzyme activity sites, similar to Figure 3B, for multiple samples. NOMe and DNase I samples were produced within DEEP (EGAO00000000236). The ATAC samples were generated by Blueprint (GSE87218) (71) and the corresponding WGBS signal tracks were attained from the IHEC epigenome data portal (72) (ERS1158277-ERS1158289). Each column contains one assay in order, NOMe, DNase I and ATAC. Each row display the measures helix turn (HelT), minor groove width (MGW), propeller twist (ProT), roll and CpG methylation bias. The shift in baseline of the methylation bias mirrors the average methylation of each sample at accessible regions.

**Figure S7.**
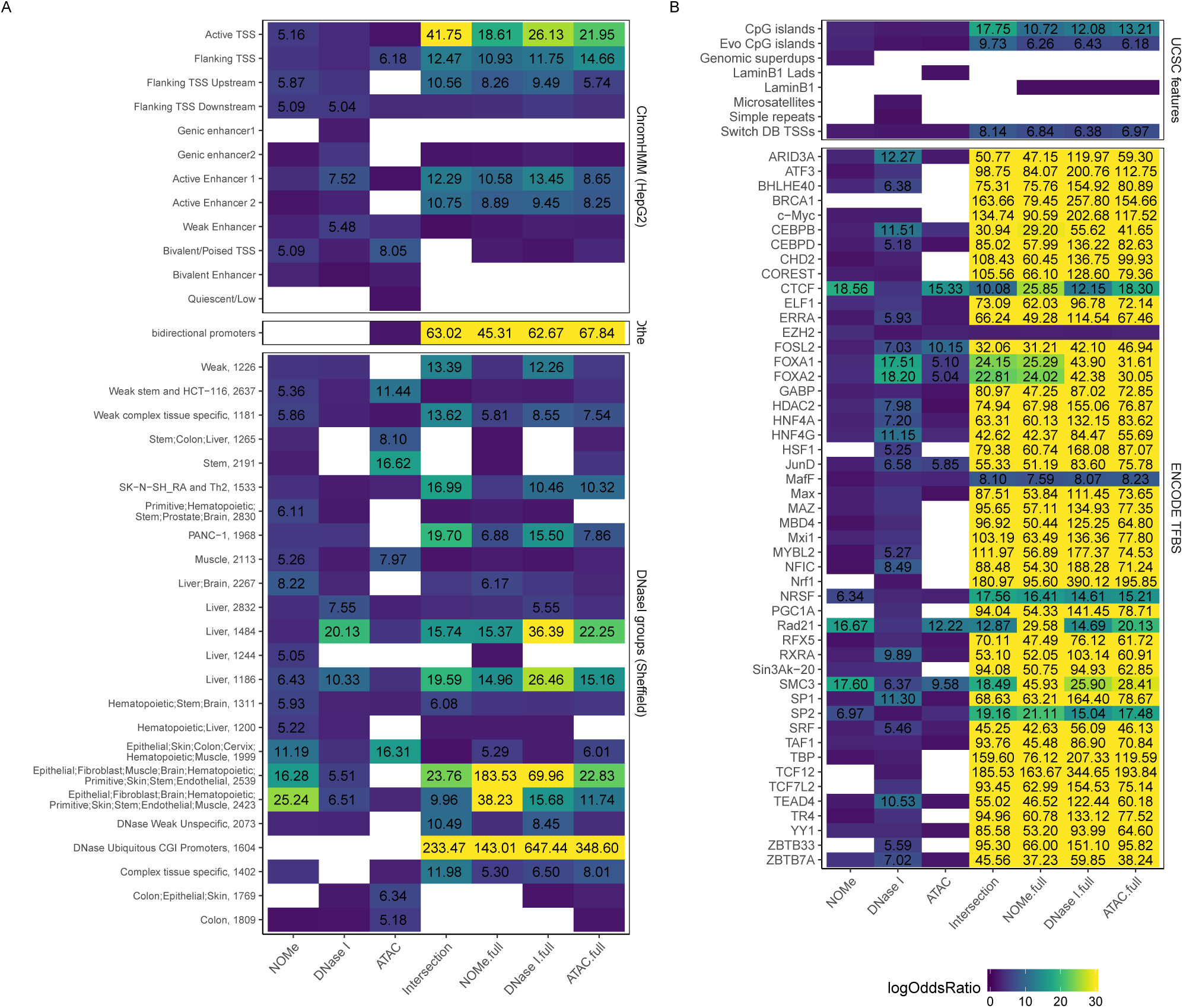
Summary of log odds ratios from a LOLA analysis of assay unique NDRs and the intersection between all three assays (see Figure 4A). Each column shows one subset. Each tile is only colored if significant (q-value¡=0.01), the color scale is cut at log odds ratios of thirty and all values above five are overlaid on the heatmap. Only TFBS and DNase I subsets with at least thousand members are displayed. A) Enrichment against ChromHMM seqmentation, bidirectional promoters and subsets of DNase I NDRs characterized as described in (73). The number suffixing the name signifies the specific DNase I subset. B) Enrichment against genomic features from UCSC and HepG2 TFBSs from ENCODE.

**Figure S8.**
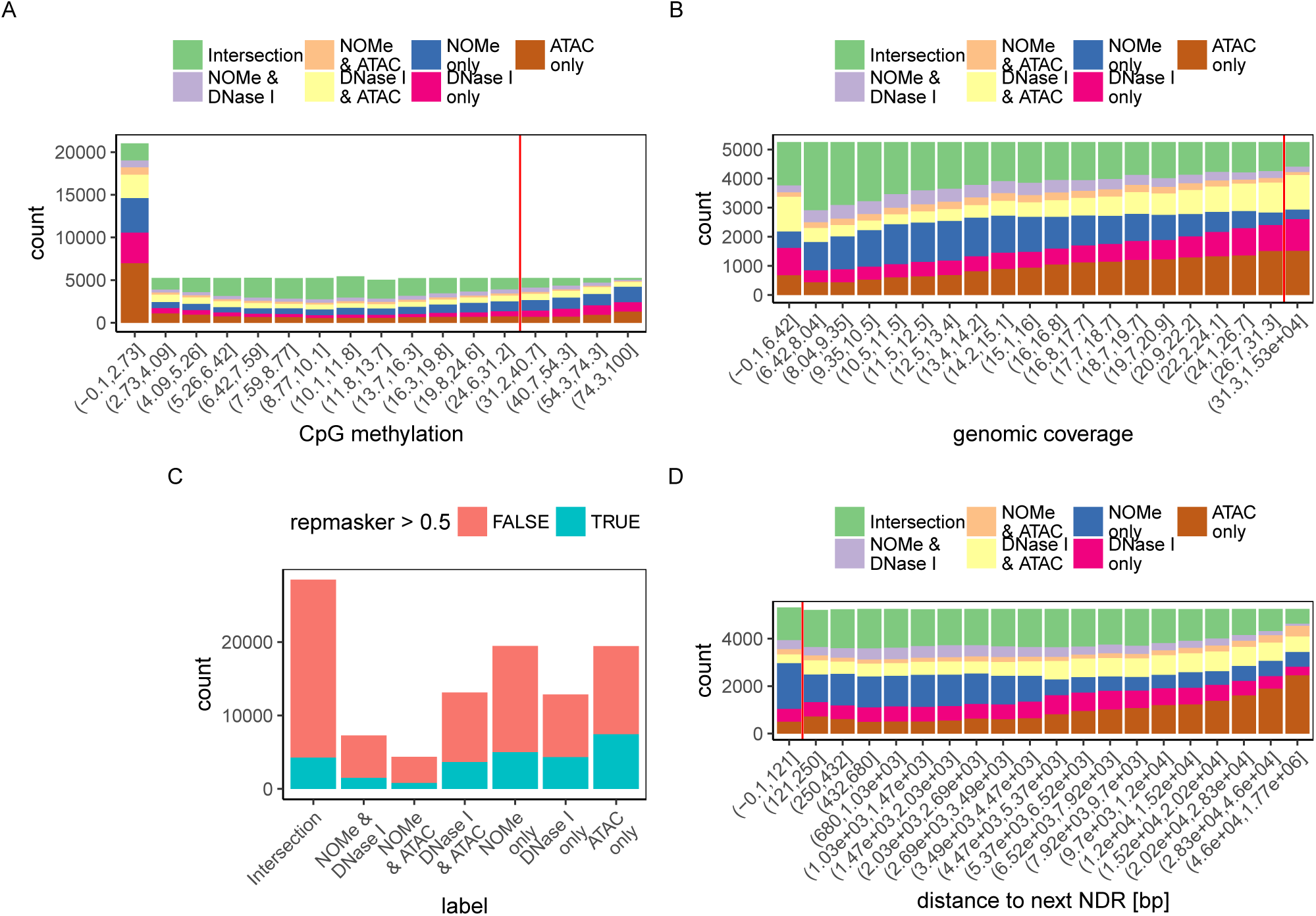
Summary of various features separating the assays. A) Average CpG methylation per NDR binned into 5% quantiles. The four smallest bins were merged as more than 15% of the NDRs had 0% CpG methylation. B) Genomic coverage, based on NOMe, split into 5% quantiles. C) Overlap with UCSC RepMasker repeats track. An NDR was considered repetitive if more than 50% of the region overlapped. D) Distance to next NDR split into 5% quantiles

**Figure S9.**
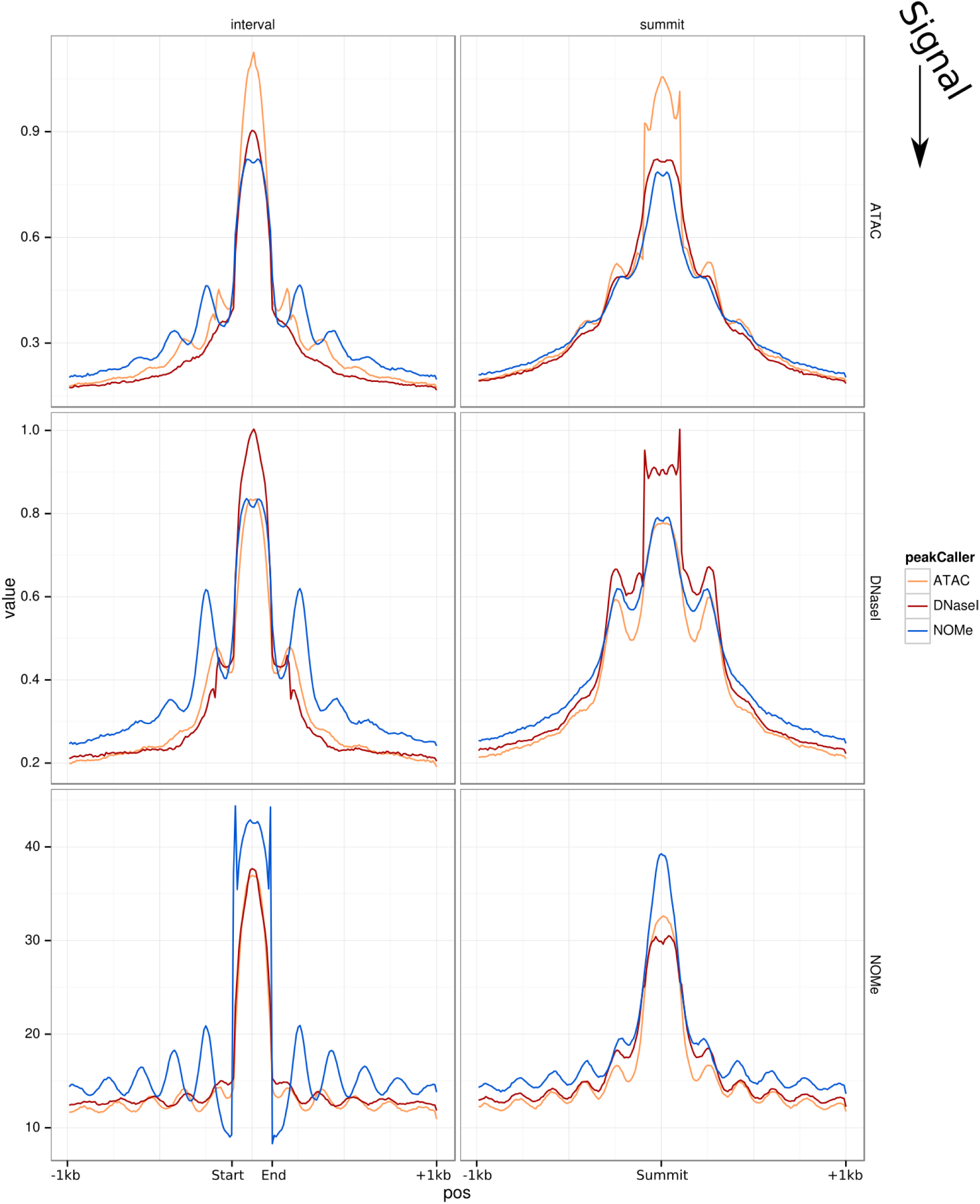
Visibility of phased signal is dependent on reference point and signal. Each panel contains the average signal in relation to a reference point. The reference point is chosen based on NDRs called in each assay; which assay is signified by the color of the line. There is one column for each type of reference point; interval) start and end of the NDR is fixed and the length between is scaled to 20 bins, summit) the peak of the NDR is fixed. Each row displays how the signal of a particular assay behaves in relation to the selected reference point. Outside the NDR, the signal is averaged in 10bp bins. Inside the NDR, this is also the case with the summit as reference point. With the NDR borders fixed, the NDR internal bin size is proportional to the NDR length.

**Figure S10.**
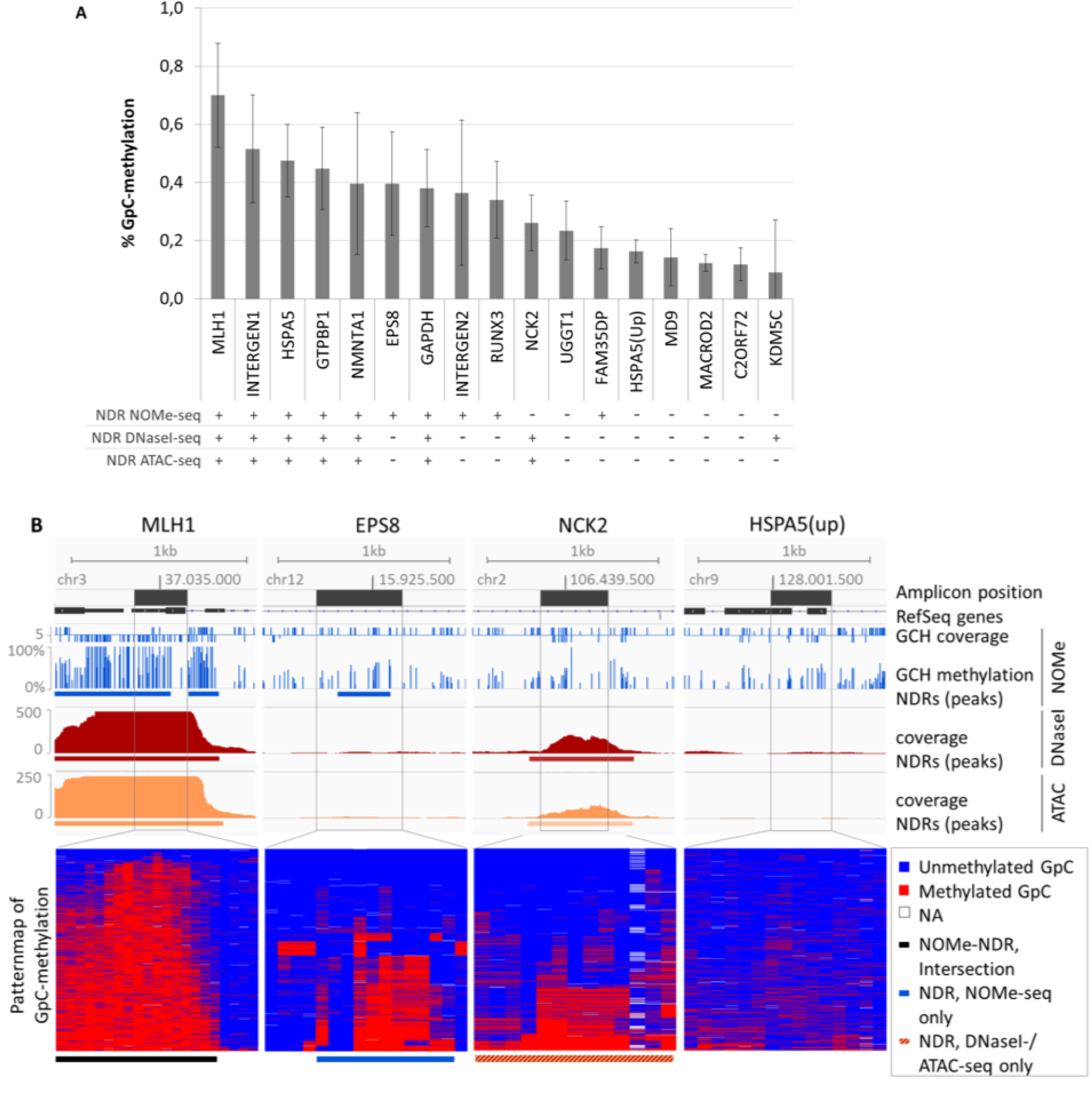
Results of targeted deep sequencing for the validation of NOMe NDRs. NDRs of Intersection, NDRs unique to NOMe-seq or DNase I/ATAC-seq as well as non-NDR regions have been analyzed by targeted deep sequencing in NOMe-samples (HepG2). A. Bar chart graphing the median GpC-methylation with standard-deviation across the regions of interest. Only GCH positions exactly overlapping with the respective NDR were used for calculations. The scheme below indicates the presence (+) or absence (-) of a NDR per Amplicon in NOMe-, DNase I-and ATAC-seq data. B. Four examples with genome-wide NOMe-, DNase I-and ATAC-profiles across regions of interest (above) and GCH-methylation pattern-maps from targeted deep sequencing for the same regions (below). Individual GpCs: columns, individual sequence-reads: rows. The overlap of the amplicon with the NDR is depicted with a bar.

**Figure S11.**
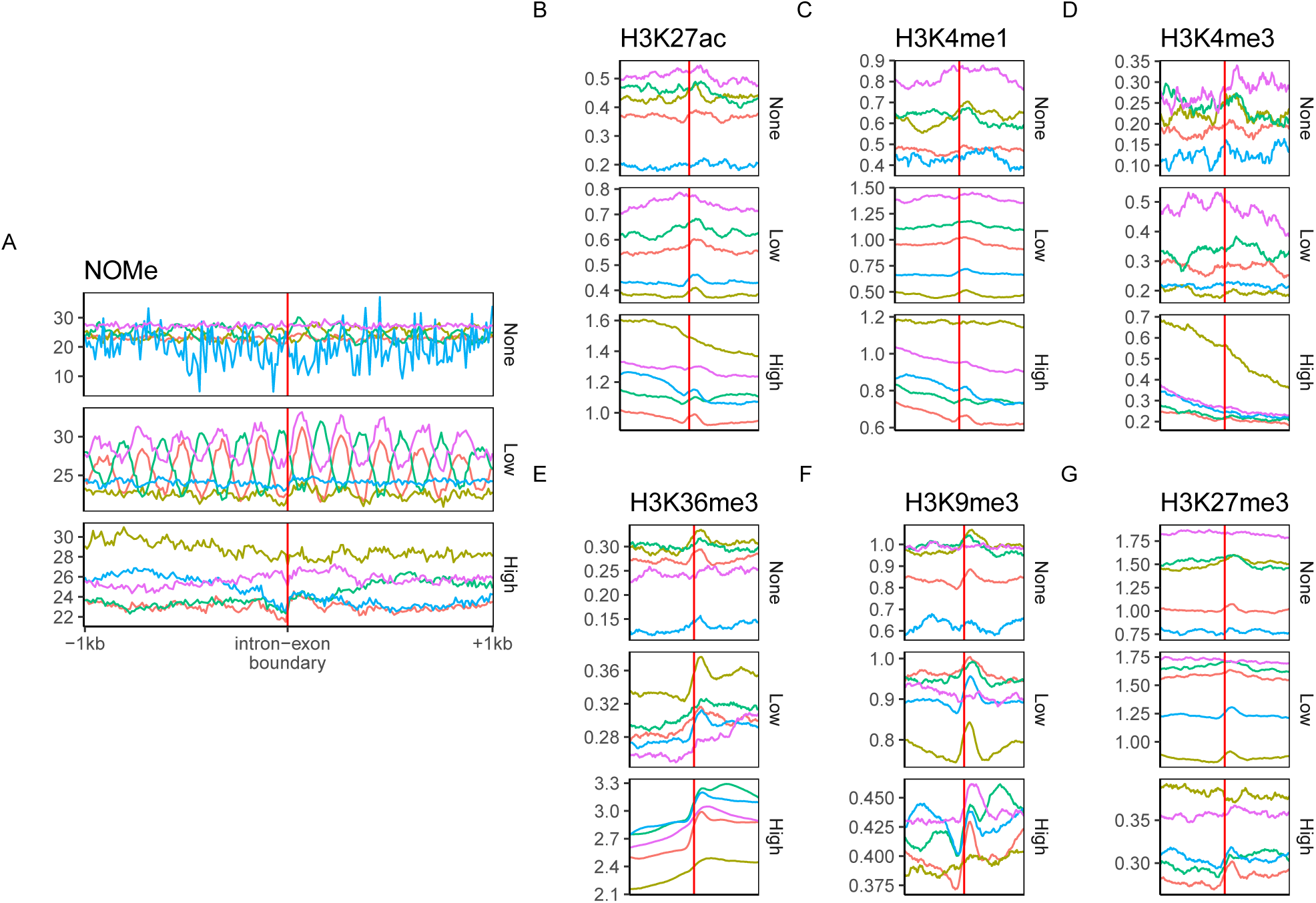
Phasing around intron-exon boundaries. The figure display aggregated NOMe and histone modification signals in 2*kb* windows centered on intron-exon boundaries. In order to avoid effects from TSS or TES, exons among the two first or last exons in a transcript or exons overlapping such exons were excluded. Signal strength was summarized in 10*bp* windows with deepTools. Exons were stratified based on the expression level of the parent gene into not expressed, low and high expressed. These groups was subsequently clustered into five clusters based on NOMe signal with K-means clustering in R. Each panel displays the average signal of each cluster with respect to different marks. The color is conserved within expression groups between marks, that is the red line in the low expressed panels are always based on the same set of boundaries.

**Figure S12.**
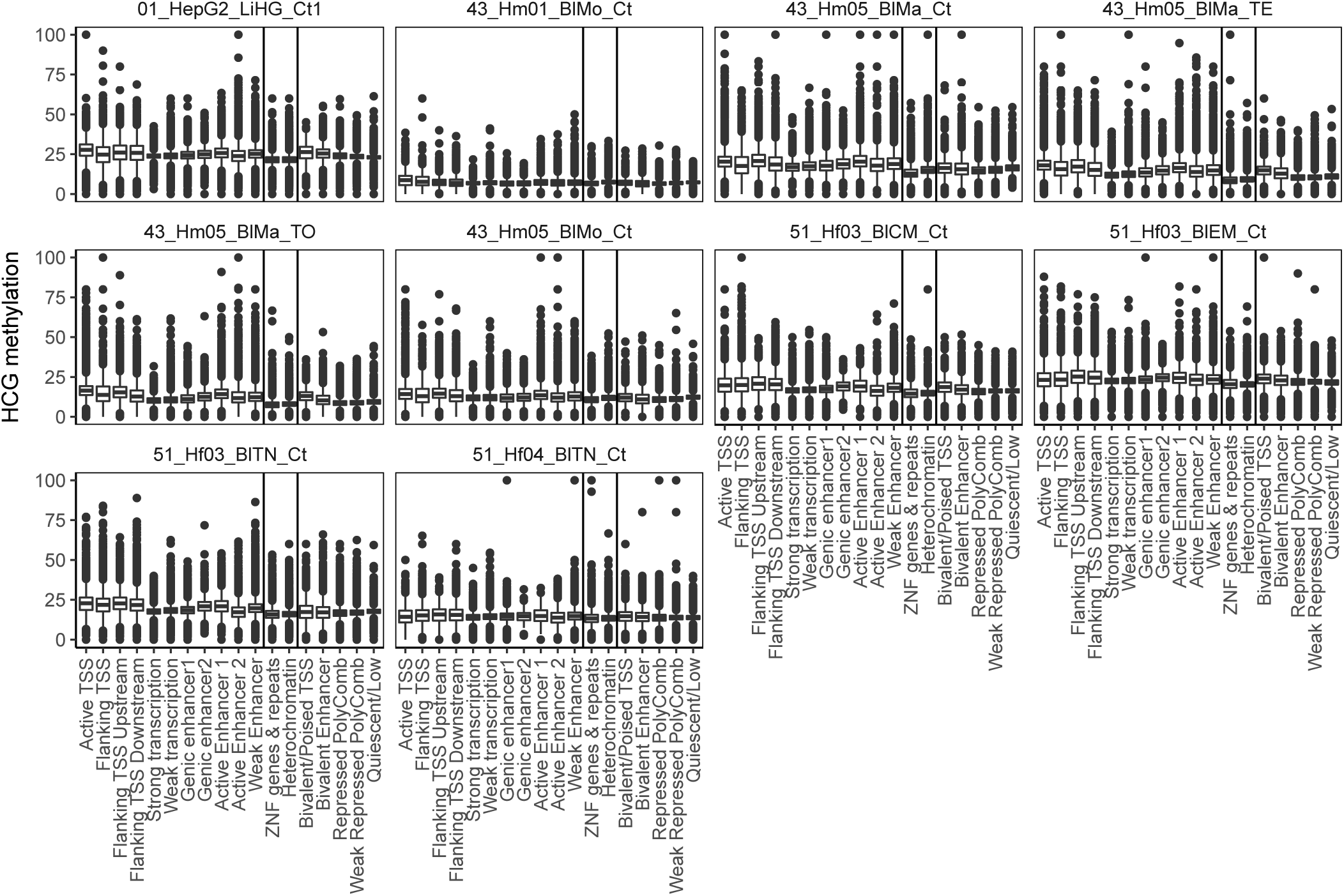
Boxplots of average NOMe GpC methylation values in genomic regions (y-axis) in ten experiments stratified over the corresponding ChromHMM segmentation states (x-axis). NDR regions are excluded.

**Figure S13.**
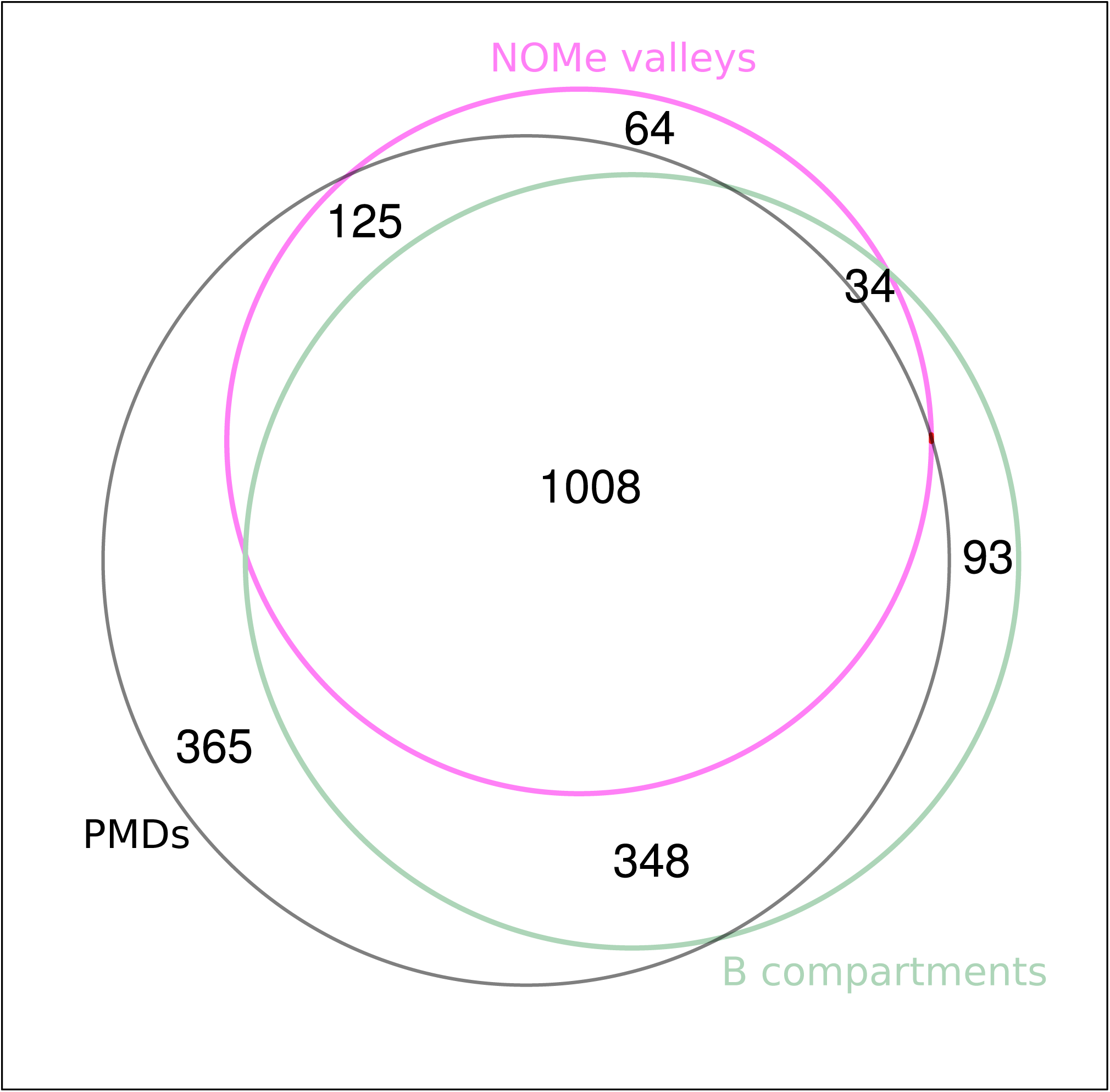
This Venn diagram displays the overlap in Mbp between NOMe valleys, Hi-C B-comparments and PMDs

**Figure S14.**
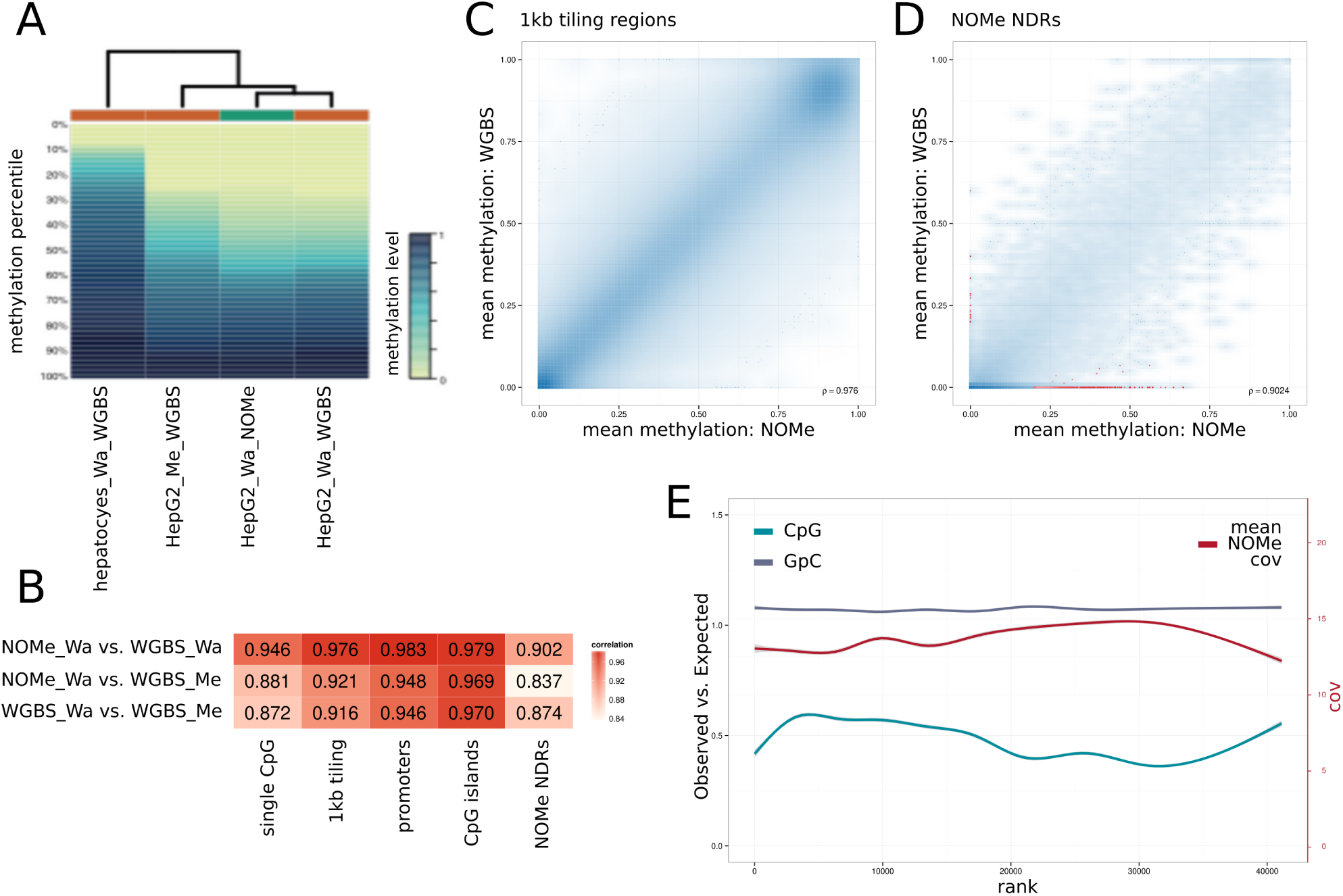
A) Distribution of CpG methylation levels in primary hepatocytes and HepG2 cells. Method and laboratory of origin are denoted in the sample suffix. Sequencing libraries for samples HepG2 Wa WGBS and HepG2 Wa NOMe were prepared from the same cell culture. Colors indicate methylation levels at percentiles. The dendrogram shows hierarchical clustering using Manhattan distance and complete linkage. B) Pearson correlation coefficients for CpG methylation profiles for HepG2 cells based on methylation levels for individual CpGs and mean methylation levels for genome-wide tiling regions, ENSEMBL gene promoters, CpG islands and regions of open chromatin as identified by NOMe-seq. C,D) Scatter plot showing mean methylation levels in HepG2 cells measured by NOMe-seq compared to the mean methylation levels of HepG2 cells measured by WGBS (averaged between the two samples characterized in A and B) across genome-wide tiling regions (1kb resolution) (C) and regions of open chromatin as identified by NOMe-seq (D). Point density is depicted by color intensity. In D) the 1000 highest-ranking differentially methylated regions (see Methods) are marked as red points. E) Observed-vs-Expected ratios for the CpG and GpC dinucleotides in the NOMe NDRs. Regions are sorted by their rank of differential methylation and generalized additive models with cubic spline smoothing were used to obtain smoothed estimates for each rank. A corresponding scatter plot smoother for mean CpG read coverage in the HepG2 NOMe sample is shown in red.

**Figure S15.**
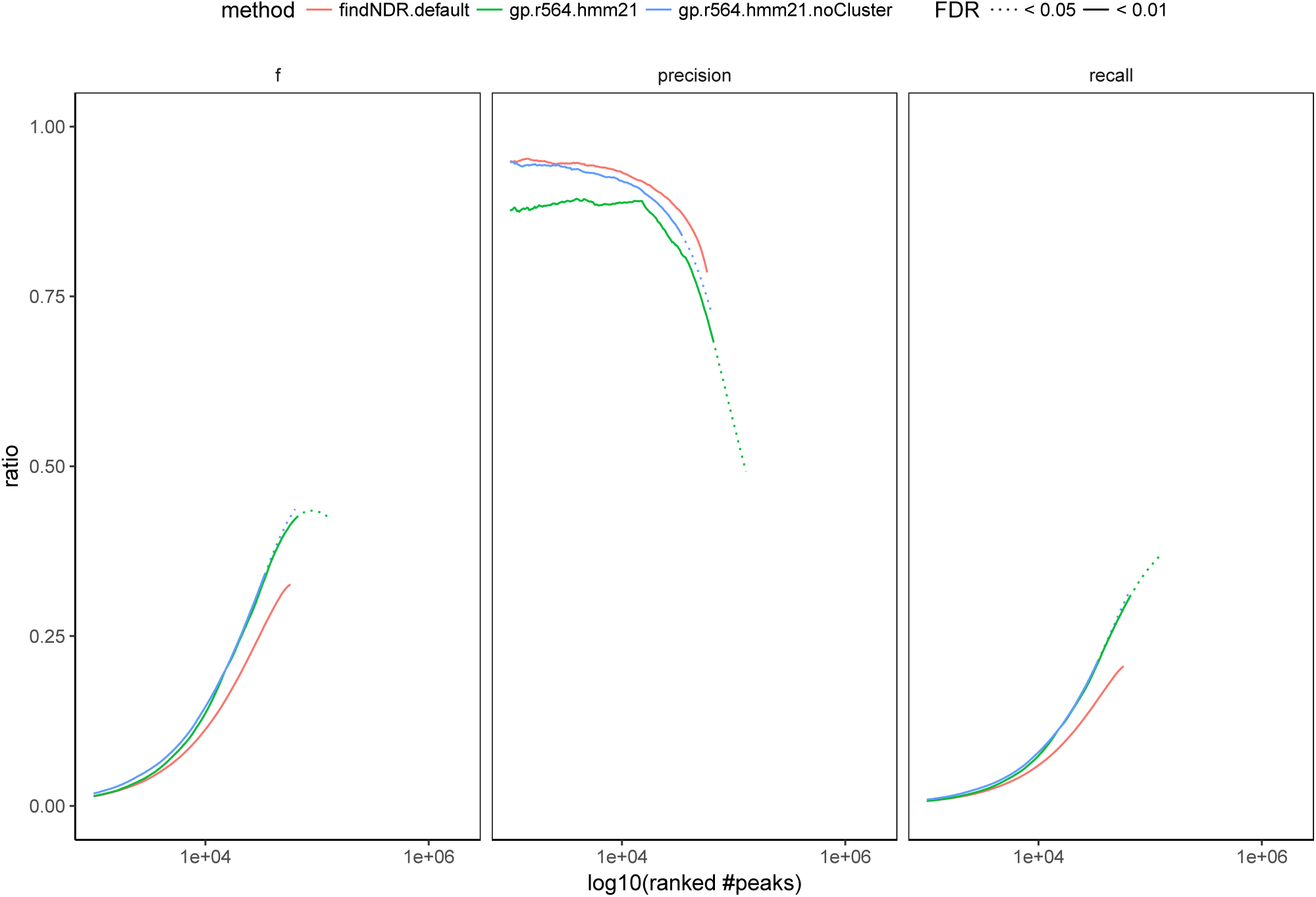
Summarizes gNOMeHMM’s ability to recapture DNase I and ATAC NDRs. The three panels show f-score, precision and recall rates for gNOMeHMM with two settings and the external software findNDR as reference in relation to the union of ATAC and DNase I NDRs. gNOMeHMM is tested with default settings and without the coverage based grouping/clustering.

## SUPPLEMENTARY TABLES

**Table S1.** Summary of HepG2 open chromatin data sets

| Assay | NOMe | DNase I | ATAC |
| --- | --- | --- | --- |
| **Cell type** | ← HepG2 → |  |  |
| **Sequencing** | ← 2x101bp → |  |  |
| **#read pairs** | 339M | 88M | 60M |
| **Duplication rate** | 2.0% | 30.8% | 58.1% |
| **Median insert size [bp]** | 267 | 160 | 206 |
| **Bisulfite conversion** | 99.7% | ← N/A → |  |
| **#peaks** | 65,683 | 62,365 | 67,675 |
| **FRiP** | N/A | 23.4% | 24.9% |

**Table S2.**
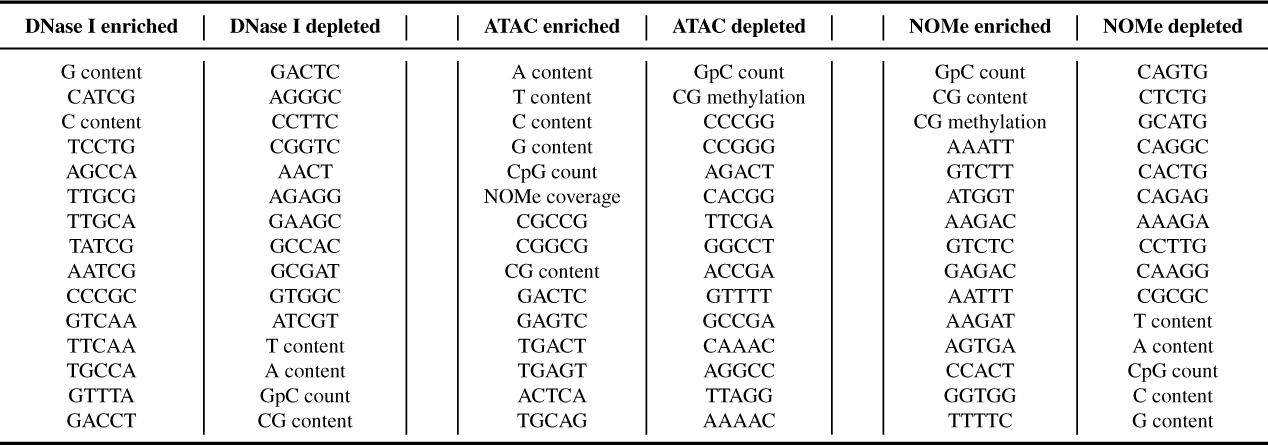
Top 15 discriminatory features for each assay. We obtained the top 15 enriched and depleted features for DNase I, ATAC and NOMe assay. The ranking of the features was obtained from the weights of the logistic regression classifier for each class (assay).

